# From letters to composed concepts: A magnetoencephalography study of reading

**DOI:** 10.1101/2020.12.07.414656

**Authors:** Graham Flick, Osama Abdullah, Liina Pylkkänen

## Abstract

Language comprehension requires the recognition of individual words and the combination of their meanings to yield complex concepts or interpretations. Rather than simple concatenation, this combinatory process often requires the insertion of unstated semantic material between words, based on thematic or feature knowledge of the concepts. For example, the phrase *horse barn* is not interpreted as a blend of a horse and a barn, but specifically a barn in which horses are kept. Mounting evidence suggests two cortical semantic hubs, in left temporoparietal and anterior temporal cortex, underpin thematic and feature concept knowledge, but much remains unclear about how these putative hubs contribute to combinatory language processing. Using magnetoencephalography, we contrasted source-localized responses to modifier-noun phrases involving thematic relations vs. feature modifications, while also examining how lower-level orthographic processing fed into responses supporting word combination. Twenty-eight participants completed three procedures examining responses to letter-strings, adjective-noun phrases, and noun-noun combinations that varied the semantic relations between words. We found that while color + noun phrases engaged the left anterior temporal lobe (150-300 ms after phrasal head), posterior temporal lobe (150-300 ms), and angular gyrus (300-450 ms), only left posterior temporal lobe responses were sensitive to implicit thematic relations between composing nouns (150-300 ms). We additionally identified a left temporo-occipital progression from orthographic to lexical processing, feeding into ventral anterior areas engaged in the combination of word meanings. Finally, by examining source signal leakage, we characterized the degree to which these responses could be distinguished from one another, using linear source estimation.

## Introduction

Successful language comprehension relies on the recognition of familiar words and the combination of their meanings to yield complex concepts or interpretations (i.e., the meaning of a phrase of sentence). However, rather than simply concatenating individual meanings, the combinatorial process often requires the specification of semantic relations between words to yield their full interpretation. For example, with little effort most English speakers will understand the phrase *horse barn* as a barn in which horses are kept, rather than a vague blend of a horse and a barn or a barn that looks like a horse. Similarly, a *trophy cabinet* is a cabinet in which trophies are kept, paralleling *horse barn* in its semantic relation, while a *metal cabinet* differs from the two, likely understood as a cabinet made of metal.

Depending on the words being combined, identifying these relations relies on different types of semantic knowledge in memory, including feature knowledge of the constituent concepts (e.g., size or color) and thematic knowledge of how they interact (Estes, 2003; Wisniewski, 1996; Wisniewski & Love 1998). While behavioral and neuroscientific investigations have demonstrated that individual words activate these aspects of conceptual knowledge in memory (e.g., Kalenine, Mirman, Middleton, & Buxbaum, 2012; Mirman & Graziano, 2012a, 2012b), comparatively little is known about how we identify and process the different semantic relations between words when they are needed to interpret multi-word concepts. Here, we used magnetoencephalography (MEG) and source localization methods to provide a detailed spatiotemporal characterization of these processes in the comprehension of two-word phrases, beginning at early stages of individual visual word recognition through to the integration of word meanings to build more complex concepts.

### Adjectives, nouns, and semantic hubs

Noun-noun combinations represent one of the most common contexts in which comprehenders must insert unstated semantic relations between words to spell out the composed meaning (Gagné & Shoben, 1997; Murphy, 1990). Previous behavioral work on these combinations has identified two ways that the constituent nouns often relate to one another (Wisniewski, 1996; Wisniewski & Love, 1998): (i) The transfer of a feature or attribute from the modifier to the head noun (attributive interpretations: *robin snake* is a red-bellied snake, where red-belly is transferred from robin) and (ii) the insertion of an implicit thematic relation between words (relational interpretations: *robin snake* is a snake that hunts robins). A recent functional magnetic resonance imaging (fMRI) study revealed separable impacts of the two interpretation types, implicating bilateral sections of temporoparietal cortex (TPC) in the processing of relation-based compounds and the left anterior temporal lobe (ATL) in the processing of attributive or feature-based compounds (Boylan, Trueswell, & Thompson-Schill, 2017). On the basis of anatomical connectivity and patterns of task-related activations, both of these regions have been suggested to house and/or function as so-called semantic hub areas (Lambon Ralph et al., 2017; Patterson, Nestor, & Rogers, 2007; Schwartz et al., 2011); a characterization reinforced by their position along gradients of resting state functional connectivity (Margulies et al., 2016). Extant accounts differ, however, in their proposals regarding what types of semantic processing are housed in one or both of these regions.

In particular, the predictions made by dual hub models of semantic knowledge (de Zubicaray, Hansen, & McMahon, 2013; Schwartz et al., 2011) are consistent with the pattern found in the results of Boylan et al., (2017). Based partly on patterns of naming errors in neuropsychological patients (Schwartz et al., 2011), these accounts propose that left posterior temporal and/or temporoparietal cortex is primarily involved in the representation of thematic information, while left anterior temporal cortex is primarily involved in representation of feature or taxonomic information (see also Mirman & Graziano, 2012a, 2012b, for supporting findings). Although conflicting results exist (Mirman, Landrigran, & Britt 2017), such accounts are bolstered by the findings that TPC and neighboring sections of the left posterior temporal lobe (PTL), particularly the posterior middle and superior temporal gyri (pMTG and pSTG), are involved in the processing of word meanings believed to rely, to a greater degree, on thematic knowledge (e.g., verbs relative to nouns; Bedny, Caramazza, Grossman, Pascual-Leone, & Saxe, 2008; Bedny, Dravida, & Saxe, 2014; Bedny & Thompson-Schill, 2006; Davis, Meunier, & Marslen-Wilson, 2004; Kable, Kan, Wilson, Thompson-Schill, & Chatterjee, 2005; Kable, Lease-Spellmeyer, & Chatterjee, 2002; Martin et al., 1995; Yu, Bi, Han, Zhu, & Law, 2012; Yu, Law, Han, Zhu, & Bi, 2011), as well as tool usage (Weisberg, van Turennout, & Martin, 2007), and action knowledge (Kalenine & Buxbaum, 2016; Kemmerer, Rudrauf, Manzel, & Tranel, 2012; see also Kalenine et al., 2009). Complementary work has highlighted the importance of the left ATL to feature knowledge and identification of objects (Baron, Thompson-Schill, Weber, & Osherson, 2010; Clarke, Taylor, & Tyler, 2011; Coutanche & Thompson-Schill, 2015; Moss et al., 2005; Tyler et al., 2004), congruent with the expected properties of a taxonomic or feature-oriented semantic hub.

Alternatively, the Controlled Semantic Cognition (CSC) framework (Jefferies et al., 2020; Lambon Ralph, Jefferies, Patterson, & Rogers, 2017) posits that the left ATL functions as the primary conceptual hub and its interaction with other areas underpins both thematic and taxonomic knowledge. Based on patterns of connectivity (Davey et al., 2016) and the results of disruption (Hallam et al., 2018; Whitney et al., 2011), as well as responses to semantic retrieval and control tasks (Davey et al., 2015; Teige et al., 2018, 2019), the CSC framework additionally proposes a finer distinction regarding the contributions of left posterior temporal and temporoparietal regions to semantic cognition. The angular gyrus (AG) is argued to support automatic semantic retrieval (i.e., straightforward semantic associations) while the pMTG is proposed to function as part of the brain’s semantic control network, supporting the flexible retrieval of contextually relevant, but non-dominant aspects of knowledge from memory (Jefferies et al., 2020). The consistent finding of activation in this posterior temporal lobe region in response to verbs relative to nouns (as well as increased responses to actions) is then postulated to be due to greater semantic control demands associated with retrieving these types of meanings from memory (Jefferies et al., 2020; Thompson et al., 2017). The proposed pMTG and AG dissociation is further supported by the results of multiple meta-analyses, which found that the left AG is reliably activated by “automatic” semantic tasks (Humphreys & Lambon Ralph, 2015), and the left pMTG by demands related to semantic control (Jackson, 2020; Noonan et al., 2013).

Assuming appropriately balanced stimuli sets, with respect to semantic control demands, the results of Boylan et al. appear to suggest that dual-hub frameworks best capture how conceptual knowledge of individual word meanings are utilized to form a coherent multi-word concept in comprehension, with thematic linkages relying on the TPC, specifically the AG, and feature-based linkages relying on the left ATL. However, the stimulus set of Boylan et al. contained many novel or otherwise unfamiliar noun-noun combinations (e.g., *sponge memory*) and the nature of the procedure provided participants with time to explicitly consider each phrase’s meaning before responding. This may have caused enhanced conscious deliberation regarding the interpretation of each combination, raising the possibility that the observed TPC and ATL responses were tied to those particular conditions. In contrast, in everyday language use we frequently encounter familiar combinations of words whose complete interpretations rely on either thematic links (e.g., *horse barn*) or the modification of a single feature of a word’s meaning. A common example of the latter is the combination of color-denoting adjectives with nouns that denote relatively simple concrete objects (e.g., *brown barn*).

To the best of our knowledge, there has not yet been a direct comparison of how simple and familiar adjective-noun and noun-noun combinations, like those above, differentially tax neural processing in the proposed hub and control regions, keeping the task, paradigm, and participants constant across the conditions. In such a contrast, strict dual hub accounts would predict, as in the Boylan et al. results, that demands related to thematic linkages in relational noun-noun compounds should engage the angular gyrus and/or posterior temporal regions, while feature-based adjective-noun modifications should exclusively modulate the left ATL’s responses. Alternatively, the left ATL may show both of these sensitivities, without corresponding modulation in the posterior temporal/temporoparietal regions, consistent with models that posit a single conceptual hub underpinning both feature and thematic knowledge.

Looking to the previous literature, a number of relevant MEG studies have examined the comprehension of color + noun phrases (e.g., red boat). The findings of these studies have consistently implicated the left anterior temporal lobe in this type of feature-oriented composition (Bemis & Pylkkänen, 2011, 2012, 2013; see Pylkkänen, 2020 for review) though, less frequently, the left angular gyrus has been implicated as well (Bemis & Pylkkänen, 2013). Notably, the limited spatial resolution of MEG linear source estimation (Hauk et al., 2011; Hillebrand & Barnes, 2002) makes it unclear exactly where in the vicinity of these regions the precise contribution to adjective-noun composition may be found. Nevertheless, evidence for a role of the AG in composition, more broadly, has also been provided by the work of Price and colleagues, who, in addition to activation differences related to modifier-noun phrases, found that atrophy in the left AG was related to impairments in combinatorial processing (Price, Bonner, Peelle, & Grossman, 2015; see Price, Peelle, Bonner, Grossman, & Hamilton, 2016 for related evidence). Focusing on noun-noun combinations exclusively, Graves et al., (2010) used fMRI to compare responses to familiar phrases (e.g., lake house) and their reversals (e.g., house lake), intended as “non-meaningful” items (i.e., devoid an immediately available interpretation). Familiar phrases elicited greater activation in the right inferior parietal lobe and angular gyrus, as well as bilateral posterior cingulate and dorsomedial prefrontal cortex. On the other hand, reversed phrases, which may require greater effort to identify or rule out a semantic relation, elicited greater activation in areas including the left inferior frontal junction, bilateral intraparietal sulcus, left posterior inferior temporal gyrus, and fusiform gyrus. Taken together, the results of these studies highlight the contributions of the anterior temporal lobes and posterior temporal/temporoparietal areas in the comprehension of multi-word concepts, and further suggest the involvement of other cortical regions, which may vary with task demands (Graves et al. 2010). The specific functional contributions of each of these regions, however, remain underspecified.

### A ventral anterior hub supporting conceptual combination

Another issue relevant to neuroanatomical accounts of semantic knowledge, and the way this knowledge is used in language comprehension, concerns how neural responses related to the sensory processing of a word feed into hub regions that are proposed underpin conceptual knowledge of that word’s meaning. This is a particularly pertinent question in visual reading, as the proposed ventral location of the ATL hub (Chen et al., 2016) lays in the vicinity of occipital and temporal lobe areas known to support visual word recognition (Cohen et al., 2002; Dehaene et al., 2010; Tarkiainen et al., 1999; Taylor et al., 2019; Woolnough et al., 2020). Previous MEG studies of reading, which provide the temporal resolution to tease apart distinct processing stages in word recognition (Pylkkänen, Stringfellow, & Marantz, 2002; Tarkiainen et al., 1999), have identified a consistent sequence of responses along the left ventral occipital and temporal lobes, progressing from sensitivities to low-level visual properties to the discrimination of word vs. symbol-string stimuli in the middle and anterior fusiform and inferior temporal gyri (Gwilliams, Lewis, & Marantz, 2016; Neophytou et al., 2018; Tarkiainen et al., 1999). This progression, which occurs within the first 200 ms following word onset, has been proposed as an electrophysiological manifestation of the hierarchy of visual word responses observed in fMRI studies (e.g., Vinckier et al., 2007), feeding into left temporal lobe regions sensitive to a word’s phonology and meaning (Taylor et al., 2019, Wang, Deng, & Booth, 2019). It has also been demonstrated to include the generators of well characterized MEG evoked components, including the M130 and M170, proposed to underpin orthographic (M130) and early lexical (M170) processing (Gwilliams et al., 2016).

Complementary to these MEG studies, investigations using intracranial recordings have provided more precise neuroanatomical detail regarding the flow of visual word processing in the ventral visual stream. Chan et al. (2011) demonstrated that a first pass of activity to the ventral anterior temporal lobe, in response to visually presented words, enabled the discrimination of semantic classes (objects vs. animals) by as early as 130 ms after onset. While this was posited to be a posterior-to-anterior flow of information, Woolnough et al., (2020) recently highlighted the possibility of the reverse, finding that after left middle fusiform discriminated between words and non-words, there was a presumptive backward (anterior to posterior) pass of lexical processing. Woolnough et al., also found that the onset latency of the middle fusiform discrimination between words and non-words was sensitive to word frequency, such that lower frequency words led to later discrimination; a pattern they proposed indexed a search in one’s mental lexicon to match visual letter-strings to a meaning.

Although these accounts may differ regarding the details of information flow, the evidence to date clearly suggests the existence of mid and/or anterior inferior temporal lobe activity linking visual letter-string inputs to a conceptual/semantic representation of word meaning. Moreover, the location of this site is at least in the vicinity of the proposed ATL semantic hub, on the ventral surface of the left anterior temporal lobe (Chen et al., 2016). The previously mentioned MEG studies of adjective-noun composition (Bemis & Pylkkänen, 2011, 2012, 2013) provide indirect evidence that this area may also contribute to semantic composition. These demonstrated that sections of the lateral anterior temporal lobe showed greater responses to word combinations than word lists (and single words), modulated by conceptual-semantic properties of the composing words (Westerlund & Pylkkänen, 2015; Zhang & Pylkkänen, 2015), at approximately the same time that Gwilliams et al., (2016) found the left anterior fusiform discriminated between letters and symbol strings (150-200 ms after word onset).

Together, these findings tentatively suggest that a ventral ATL region may contribute to lexical processing and the construction of multi-word concepts by 150-200 ms after the onset of a visually presented word (e.g., a phrasal head). More specifically, we propose that the ventral ATL may house a transformation from representations that are still, to some degree, based on visual input, to conceptual-semantic representations that can then be fed into (semantic) compositional processes. If this is the case, we would expect that those anterior fusiform areas that distinguish between letter-strings and symbols (by 150 ms after onset, on the basis of previous findings) to also show enhanced responses to examples of conceptual combination (e.g., red + boat), with dual- and single-hub models making different predictions regarding what types of combinations would modulate responses here (thematic vs. feature, see above). This remains an open question, however, as there has yet to be a study that jointly localizes the temporally resolved ventral stream responses to letter and word stimuli, as done by Tarkiainen et al., (1999) and Gwilliams et al., (2016), as well as the left anterior temporal lobe’s responses to word combinations.

### The Present Study

Here, we conducted a series of MEG studies in a common sample of participants to address each of the issues outlined above. Participants completed three procedures that examined early visual processing through to conceptual combination. Specifically, these probed the processing of (i) visually presented letter-string and word stimuli; (ii) combinations of color adjectives and nouns compared to individual nouns and two-word lists; and (iii), combinations of nouns that systematically varied the semantic relation between constituents. The first two procedures were replications of previous studies (Gwilliams et al., 2016 and Bemis & Pylkkänen, 2011, respectively). The third manipulation consisted of primarily familiar noun-noun combinations that shared a common head but modified either visual/material properties (e.g., *metal cabinet* is a cabinet made of metal) or a spatial/functional relation with another noun (e.g., *trophy cabinet* is a cabinet that holds trophies). The former represents a class of non-predicating noun-noun compounds, which can be adequately, though not perfectly, paraphrased without specification of a thematic relation (i.e., “a metal cabinet is a cabinet that is metal”), while the latter require complete specification of a thematic relation (i.e., the phrase “trophy cabinet” does not mean “a cabinet that is trophies”). By also including the same head noun modified by a color-denoting adjective (e.g., green cabinet), this allowed us to identify neural responses modulated by the specific type of relation between the two composing nouns (thematic vs. non-thematic) and assess how these responses patterned across the more general contrast of noun-noun vs. adjective-noun combinations.

Using linear source estimation, we examined responses in three theoretically relevant regions of interest (the left anterior temporal lobe, angular gyrus, and posterior temporal lobe), based on the proposals of dual and single hub neuroanatomical models. By also localizing responses related to visual word and letter recognition (replicating Gwilliams et al., 2016), we positioned these effects with respect to the ventral stream’s responses to visual words and identified the earliest point along this stream that was sensitive to composition. Importantly, using MEG to examine responses in these adjacent regions required careful consideration of the limitations of source estimation methods, which tend to blur together responses in neighboring cortical areas, weakening conclusions one can make about the localization and independence of source-level patterns (Hauk et al., 2011; Hauk et al., 2019; Liu et al., 2002; Molins et al., 2008). Assessing whether this is the case in the relevant regions was thus an important methodological step toward understanding how MEG source modelling may be appropriately applied to these problems. For this reason, we quantified the source-level blurring and estimated localization accuracy in each region of interest (ROI). Altogether, this provided a detailed characterization of how the brain’s reading and semantic systems support incremental language comprehension, as well as the limitations of examining these systems with MEG.

## Materials and Methods

### Participants

Twenty-eight right-handed native English speakers (20 females, 8 males, mean age = 28.14 years, sd = 10.25 years) took part in the study. All participants had healthy or corrected-to-healthy vision and healthy hearing. Twenty-one of the participants also took part in an MRI experiment that included collection of a high-resolution anatomical MRI, which was used in the source estimation of MEG responses.

### Stimuli & Experimental Design

The complete experimental design consisted of three separate MEG experiments to probe the processing of (1) visual letter-strings and words, (2) adjective-noun composition; and (3) noun-noun composition. Figure 1 displays example trials from each experiment. More details are provided in each of the following sections.

**Figure 1.**
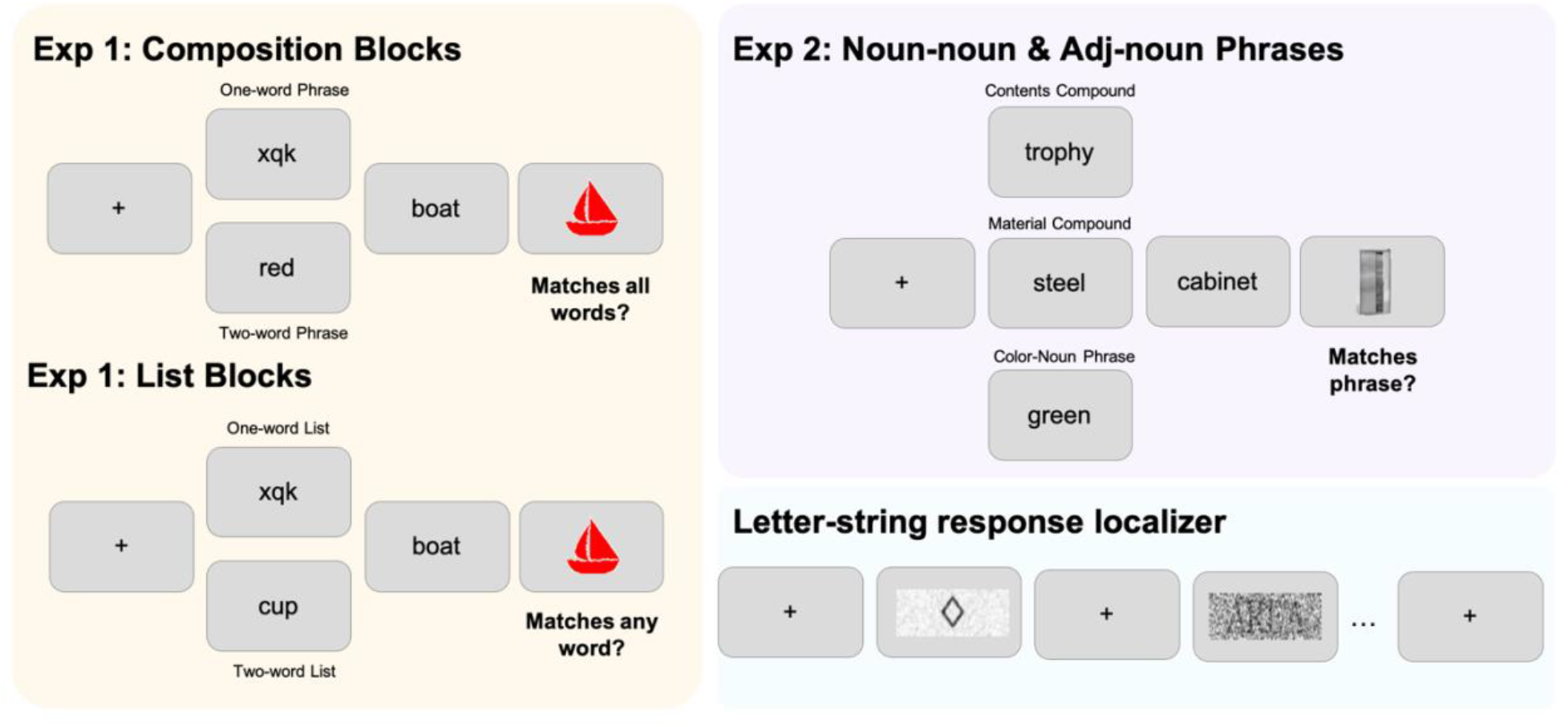
Trial structures for each experimental procedure. Left: Experiment 1 replicated the procedure of Bemis & Pylkkänen (2011) and contained two types of blocks. In Composition Blocks, participants read two-word phrases (red boat) or one-word phrases, wherein the modifier was replaced with an unpronounceable consonant string (xqk boat). Their task was to indicate if a subsequent picture matched all of the words they read on that trial. In List Blocks (bottom), participants read lists of two nouns (cup, boat) or one-word lists (xqk, boat), and then indicated if a subsequent picture matched any of the words on that trial. Top-right: In Experiment 2, designed to parallel Experiment 1, participants read three types of phrasal stimuli: Contents Compounds (modifier specifies the function/contents of the head noun, considered a thematic relationship; trophy cabinet), Material Compounds (modifier specifies the material of the head noun, considered a non-thematic relationship; metal cabinet), or adjective-noun phrases in which the adjective denoted a color (green cabinet; non-thematic). They were then asked to judge whether a following picture matched the meaning of the phrase they read on each trial. Bottom-right: Participants also completed a letter-string response localizer task (see Gwilliams et al., 2016 for full details), which involved passive viewing of letter- and symbol-string stimuli. See the main text for complete details on all three procedures.

#### Letter-string response localizer

First, to characterise responses to visual letter-strings, we adopted the localiser of Gwilliams et al., (2016, see original paper for full details), developed from the work of Tarkiaenen et al., (1999). Briefly, participants passively viewed four types of visual stimuli: An individual letter (e.g., A), a combination of four letters that formed a familiar, disyllabic word (e.g., ATOM), a single shape symbol length-matched to the one-letter stimuli (e.g., a single square), and four shapes length matched to the four-letter word stimuli (e.g., a square, circle, triangle, and diamond; see Figure 2 for example stimuli). Letter and word stimuli were embedded in two levels of visual noise, defined as zero-mean Gaussian distributions with variances of 0.0234 (low) and 1.5 (high). Altogether, this facilitated the isolation of neural responses sensitive to low-level visual properties of letter-strings (e.g., high vs. low visual noise across letters and words) and differences in stimulus type (e.g., letters and words vs. length-matched symbol strings). Notably, the four-letter stimuli in this localizer were pronounceable words, rather than pseudowords, meaning that their contrast with length-matched symbol strings would reveal responses that may be plausibly related to orthography, phonology, or word meaning.

**Figure 2.**
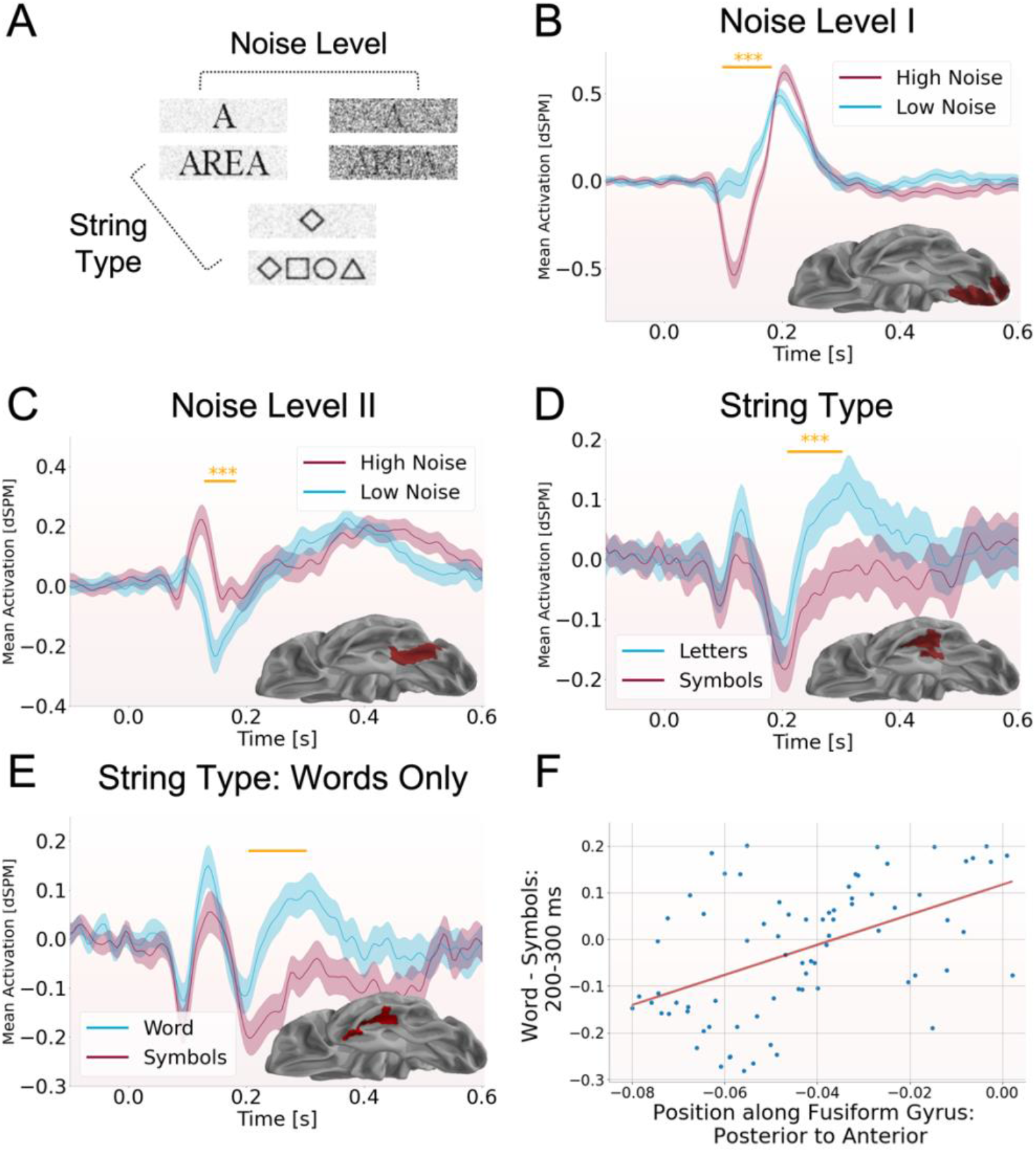
Letter-string response localizer. (A) Participants viewed one-letter items embedded in high and low visual noise, four-letter words embedded in high and low visual noise, and one- and four-unit length symbol strings, in low visual noise. (B) In left occipital cortex, high noise letter-string stimuli elicited a negative polarity response component between 100 and 180 ms post stimulus onset that was absent in responses to the clean letter-string stimuli (p < 0.0001). (C) In the left posterior fusiform gyrus, high and low noise letter-string stimuli elicited opposite polarity responses between 130 and 180 ms (p < 0.0001). (D) Responses in the left anterior fusiform and inferior temporal gyrus showed a dissociation in responses to letter-string stimuli and symbol stimuli (across both 1- and 4-unit lengths), with greater positive amplitudes elicited by letter-strings than their symbol counterparts, between 210 and 300 ms after stimulus onset (p < 0.0005). (E) When the same String Type analysis was performed on only the four-unit stimuli (words and four symbols) a more anterior fusiform cluster was found (205-300 ms, p < 0.008) showing the same response dissociation. (F) A regression analysis of posterior-to-anterior position along the left fusiform gyrus and the difference between word and four-unit symbol strings identified a significant positive relationship, revealing that the difference between the two increased along this axis. This relationship was not found when comparing individual letters and symbols.

#### Experiment 1: adjective-noun composition

Second, to probe responses related to adjective-noun composition, as in simple color-noun phrases (e.g., red + boat), we performed a truncated replication of the MEG study conducted by Bemis & Pylkkänen (2011), using all of the original stimuli and procedures (see original paper for full details). Participants performed two tasks in a blocked fashion, with each block made up of two- and one-word trials. In “Composition” blocks, two-word trials consisted of a color-denoting adjective (e.g., red) followed by on of twenty-five concrete nouns (e.g., boat). On one-word trials the adjective was replaced by a length-matched, non-pronounceable, consonant string. Following presentation of these stimuli, an image of a colored line drawing appeared on screen and participants indicated whether it matched or mismatched all of the words on that trial (i.e., was it a red boat?).

In “List” blocks, two-word trials consisted of a pair of nouns (e.g., cup, boat), rather than a phrase, while one-word trials again replaced the first position item with a length-matched consonant string. As in Composition blocks, an image appeared following the word stimuli, but here participants were asked to indicate whether the pictured matched any of the words on that trial (i.e., was it either a cup or boat?). On one-word trials, in each task, participants only needed to consider the single noun they encountered. Each participant completed fifty trials in each condition in four alternating Composition/List blocks. This represented one half of the complete procedure of Bemis and Pylkkänen (2011; 100 trials in each condition). Stimuli lists were pseudo-randomly generated for each participant such that no unique combination of word/non-word stimuli could be repeated more than twice, no list could consist of a repeated noun, and each condition was balanced on the overall length of the first position item. We note that in the original paradigm, Bemis & Pylkkänen (2011) repeated the same set of images in each condition. Here, instead, we pseudo-randomly selected a set of 200 total images for each participant, which were then assigned to each trial to ensure that half of each condition was followed by a match and half followed by a mismatch. In the two-word conditions, the image assignments were also appropriately balanced on the relevance of the first or second item to determining whether the correct response was match or mismatch.

#### Experiment 2: Noun-noun composition

The third procedure was modeled after the previously described adjective-noun composition paradigm (particularly the two-word Composition trials), but contrasted responses to noun-noun combinations that differed in their relational structures between modifier and head, as well as the same set of head nouns modified by color-denoting adjectives (e.g., trophy cabinet vs. metal cabinet vs. green cabinet). The construction of the stimulus set began with the selection of fourteen container-denoting head nouns, which could be straightforwardly modified by a preceding noun specifying the material they are made of (e.g., metal cabinet) as well as a noun that in a spatial/functional relationship with the head (e.g., a trophy cabinet is a cabinet that functions by containing trophies). For convenience, we refer to the former as Material modifiers (and their corresponding whole phrases as Material Compounds) and the latter as Contents modifiers (and Contents Compounds). Fourteen Content modifiers were selected, and each was paired with between 2 and 5 head nouns to yield a set of 42 items (see Supplementary Materials for the complete stimulus set). Fourteen Material modifiers were then selected such that each was assigned to replace one Contents modifier in all of its phrases, creating a set of 42 Material Compounds. Notably, although a subset of the phrases in both conditions were novel (e.g., shampoo cup), the complete set in each condition was designed to include primarily familiar noun-noun combinations (see below for more details on familiarity).

Lexical and phrasal characteristics of the stimuli were extracted from the Corpus of Contemporary American English (Davies, 2009) and the English Lexicon Project (Balota et al., 2007). The selected modifiers were appropriately balanced on word length (Contents: mean = 6.00, sd = 1.82; Material: mean = 5.69, sd = 1.73), lexical frequency (Contents: mean = 13213.98, sd = 9592.3; Material: mean = 13816.86, sd = 10279.7), and mean reaction time in lexical decision tasks, as reported in the English Lexicon Project (Contents: mean = 635.53 ms, sd = 65.48 ms; Material: mean = 638.56, sd = 78.24 ms). The two sets of compounds were balanced on bigram frequency when the head noun was marked as a singular noun (Contents: mean = 41.00, sd = 81.66; Material: mean = 37.67, sd = 74.17) and transition probability from modifier to head (Contents: mean = 0.004, sd = 0.009; Material: mean = 0.004, sd = 0.008). The two sets also had comparable numbers of phrases with zero bigram frequency in the Corpus of Contemporary American English (9 items in the Material Compounds and 13 in the Contents Compounds). For context, of the sixty-four items in each of Boylan et al.’s (2017) relational and attributive compound sets (selected from previous studies), twenty-six and thirty-seven had zero bigram counts in this corpus, respectively. We additionally conducted two stimulus norming studies on Amazon Mechanical Turk (AMT) to confirm that the sets were balanced on overall familiarity and readers’ tendencies to interpret the phrase using the intended relational structure (i.e., a metal cabinet is a cabinet made of metal, rather than a cabinet that holds metal). For complete details of these norming studies, see the Supplementary Materials. In brief, Contents and Material compounds were rated as similarly familiar (using a Likert rating scale where 1 indicates completely unfamiliar and 7 indicates extremely familiar: Material: m = 4.750, sd = 1.021; Contents: m = 4.838, sd = 1.351) and were consistently interpreted in the intended fashion.

Once the complete noun-noun stimuli had been finalized, we also matched each modifier to one of fourteen color-denoting adjectives. It was not possible to balance the noun modifiers and color adjectives on the same properties as the two original conditions, with the color adjectives being more frequent on average and showing greater variability in their frequency (m = 92030.74, sd = 105749.29), shorter in length (m = 5.02, sd = 0.92), and appearing with the head nouns with lower bigram counts (m = 22.83, sd = 55.94) and transition probabilities (m = 3.6 × 10^−4^, sd = 0.001). For this reason, all critical contrasts were conducted between the Material and Contents Compounds, and the adjective-noun stimuli in this procedure were only considered in post-hoc assessments. From one perspective, Material modifiers and Color modifiers are more similar to each other than the Color modifiers are to the Contents modifiers, as the former pair specifies simple physical/visual attributes of the head noun that often covary (e.g., materials such as leather or steel each have a prototypical color). Many material modifiers can also be licit in syntactic contexts that are typically characteristic of adjectives, such as following linking verbs (e.g., that looked painful vs. that looked metal vs. that looked trophy). Moreover, as mentioned in the Introduction, both the Material and color modifiers can be adequately paraphrased as non-predicating relations with the head noun while the Contents Compounds cannot. That is, while “a green cabinet is a cabinet that is green” is an adequate paraphrase and “a metal cabinet is a cabinet that is metal” is underspecified by still acceptable, the phrase trophy cabinet cannot be paraphrased as “a cabinet that is (a) trophy”. While only an informal analysis, this contrast serves to highly the extra thematic relation that is required in the Contents Compounds but can be acceptably dropped in the Material Compounds and color-noun phrases.

Lastly, in addition to the phrasal stimuli, we also presented all individual constituent words in the third procedure in isolation, for the purposes of analyses that are beyond the scope of this paper. This was implemented by splitting the experiment into alternating Two-word and One-word blocks. In the former, participants saw a randomly ordered set of the adjective-noun and noun-noun stimuli, while in the latter they saw a randomly ordered set of the individual nouns and adjectives. Each modifier constituent was presented in isolation an equal number of times as it was encountered in the phrasal contexts, while head nouns were each presented once.

As in the Composition trials of the adjective-noun procedure (Experiment 1), each trial was followed by an image and participants indicated whether it matched or mismatched all of the words encountered on that trial. Participants were additionally instructed that if the image contained any depiction of the words on the trial, they should respond “match”. This was specified to handle the inclusion of Material and Color adjectives in one-word blocks, as these specify only a property of objects depicted in an image (i.e., is “green” in this image?). No “list tasks” were performed in the noun-noun composition procedure. One-hundred and twenty-six images that depicted the set of adjective-noun and noun-noun phrases were selected for use as task images. A list of stimulus-image pairings was pseudo-randomly generated for each participant, such that half of the trials contained a match and half of the trials contained a mismatch, in each condition, in each block.

### Procedure

All experimental procedures took place in the Magnetoencephalography and Magnetic Resonance Imaging Laboratories of New York University Abu Dhabi. Every participant provided informed consent prior to taking part in the research and all procedures were approved by the Institutional Review Board of New York University Abu Dhabi. Prior to beginning MEG procedures, each participant had their head shape, the future locations of five head position indicator coils, and the position of three fiducial landmarks (nasion, and left and right tragi), digitally recorded using a Polhemus FastSCAN system (Polhemus, Vermont, USA). Each participant completed the MEG procedures in the following order: adjective-noun composition (Experiment 1), noun-noun composition (Experiment 2), and then the letter-string response localizer. The order of the blocks in Experiments 1 and 2 were counterbalanced across participants. Both Experiments 1 and 2 began with a brief instruction period followed by a practice session of 20 trials of each type in that experiment. The letter-string response localizer paradigm was completed exactly as described by Gwilliams et al., (2016).

Experiments 1 and 2 each used rapid serial visual presentation (RSVP) paradigms. Example trials are shown in Figure 1. Every trial in both procedures began with the presentation of a fixation cross for 300 ms followed by a blank screen for 300 ms, and then each word on that trial presented for 300 ms, with a 300 ms blank screen following its offset. Task images were presented on screen until participants pressed the button to respond. The duration of each interval between successive trials was randomly sampled from a uniform distribution consisting of discrete values of 300, 400, 500, 600, and 700 ms.

### Data Collection and Preprocessing

Continuous MEG data were acquired throughout all experimental procedures with a 208-channel Kanazawa Institute of Technology system (Eagle Technology, Japan) at a sampling rate of 1000 Hz. Online high- and low-pass filters of 0.1 and 200 Hz were used during data collection. Head position indicator coils were used to record each participant’s head position, relative to the MEG sensors, before and after each experimental procedure. Twenty-one of the participants also took part in an MRI session that included the acquisition of T1- and T2-weighted high-resolution anatomical MRIs on a 3-Tesla MAGNETOM Prisma scanner (Siemens Healthineers, Erlangen, Germany). The high-resolution anatomical scans along with associated field-maps were acquired and preprocessed according to the Human Connectome Project’s Young Adults protocols (Glasser et al., 2013). For all relevant participants, MRI acquisition followed participation in the MEG experiment by no greater than 14 days.

MEG data from each participant were first cleaned of environmental electromagnetic noise using the Continuously Adjusted Least-Squares Method (Adachi et al., 2001) based on data collected at three reference channels placed away from the head. All remaining preprocessing analysis steps were completed using the MNE-Python (v. 0.20; Gramfort et al., 2013) and Eelbrain (v. 0.28; DOI: 10.5281/zenodo.1444075) packages in the Python computing environment. The data were then low-pass filtered at 40 Hz and Independent Component Analysis was used to remove data patterns matching the profile of known artifacts (eye blinks, movement-related activity, and well-characterized external noise sources). The continuous MEG data were then split into epochs around the onset of critical events in each procedure. Baseline correction was applied using prestimulus intervals and channel noise covariance matrices were estimated for each participant from the concatenation of these intervals. In analyses of Experiments 1 and 2, trials in each condition were averaged within each participant’s data set before source estimation. In the letter-string response localizer, following the original analysis (Gwilliams et al., 2016), individual trial data were used in source estimation without averaging.

For those participants with anatomical MRIs available, images were processed with the automated segmentation algorithms of the Freesurfer software suite (http://surfer.nmr.mgh.harvard.edu/) to generate a cortical surface reconstruction and corresponding parcellations for each individual (Dale, Fischl, & Sereno, 1999; Desikan et al., 2006; Fischl. Sereno, & Dale, 1999; Fischl et al., 2004). For the remaining participants, the cortical surface of Freesurfers’s “fsaverage” template and the corresponding parcellations were scaled to match the head shape and location of fiducial landmarks from each participant. MEG and MRI coordinate spaces were co-registered based on the location of fiducial landmarks and head position indicator coils. Forward models were estimated for each participant using a single layer conductance boundary element model. The L2-minimum norm method was used to estimate source-level activity in each participant’s cortical surface, using dynamic statistical parameter mapping (dSPM) to account for superficial source bias. The regularization signal-to-noise ratio parameter was set to 3 for all analyses involving averaging in sensor space prior source estimation and 2 for the letter-string response localizer (motivated by original analyses).

As has been discussed in previous work (Dale & Sereno, 1993; Gwilliams et al., 2016; Lin et al., 2006), anatomical information concerning the geometry of each individual’s cortical surface can be used to constrain the minimum norm estimate. Specifically, the orientation of sources distributed throughout the cortical surface can be specified to lie perpendicular to it. The use of this “fixed” orientation is motivated, in large part, by the known sensitivity of MEG to postsynaptic potentials in pyramidal cells, which lie perpendicular to the cortical surface (Hamalainen et al., 1993; Okada, Wu, & Kyuhou,1997). Alternatively, the orientation of the source with respect to the cortical surface can be discarded (“free” orientation analysis) and, instead, only the magnitude (norm) of the vector returned. In the previous work from which our letter-string response localizer was taken, Gwilliams et al., (2016) demonstrated that the discarding of source orientation information (i.e., using free orientation estimates) obviated response dynamics that were relevant to the processing of visual word stimuli. This motivated our use of the fixed orientation option in the primary analysis of our data. In all source estimations, cortical patch statistics were used to define the orientation normal to the surface (Lin et al., 2006).

### Statistical Analyses of MEG Data

All analyses of data collected in the letter-string response localizer were directly motivated by the results of Gwilliams et al., (2016). These focused on the identification of three primary response components: (i) The Type I Noise component localized to lateral and ventral occipital cortex between 100 and 130 ms after stimulus onset, appearing as greater negative polarity in response to high visual noise than low visual noise; (ii) The Type II Noise component, localized to the left posterior fusiform gyrus and appearing as a polarity divergence between high and low visual noise stimuli approximately 100-130 ms after stimulus onset, and (iii), The String Type component, localized to the anterior fusiform and inferior temporal gyri and appearing as greater response amplitudes to letter-string stimuli as compared to length-matched symbols between 130 and 250 ms.

Data from each individual trial were converted to distributed source amplitude estimates using each subject’s noise-normalized minimum norm inverse solution. Once in source-space, linear regression models were fit to the data at each time point across all trials. Models included the binary-coded predictors of interest (noise type, string type) and, as a nuisance variable, the elapsed number of trials. Spatiotemporal cluster tests (see Maris & Oostenveld, 2007) were then performed on the regression coefficients for each predictor, across participants, using one-sample t-tests for significant deviations from zero. In all analyses, including those for Experiment 1 and 2, clusters were formed using the threshold-free cluster enhancement method (TFCE; Smith & Nichols, 2009). Permutation p-values were estimated by random sign flipping of the coefficients ten thousand times and repeating the clustering procedure, then comparing the observed cluster magnitude (the sum of constituent statistics in the cluster) to the resulting distribution of largest cluster magnitudes from each permutation.

All ROIs were defined using the anatomical parcellations of the fsaverage cortical surface from Freesurfer (Desikan et al., 2006; Van Essen, 2005). The tests for Noise Type I and II components were performed on activity between 100 and 180 ms after stimulus onset, in a spatiotemporal region of interest (ROI) encompassing the lateral occipital, cuneus, lingual, pericalcarine, fusiform, middle temporal, and inferior temporal gyri. The test for the String Type component was performed in only the anterior half of the ROI, between 130 and 300 ms after stimulus onset. The more constrained, anterior ROI was motivated by the localization of the component by Gwilliams et al., and the present interest in isolating a ventral anterior temporal lobe region. The longer duration test window, extending to 300 ms, was motivated by the desire to identify potential overlap in time with adjective-noun composition effects (see below).

In the analysis of the Experiment 1 data, spatiotemporal cluster tests were used to identify the interaction of interest: responses that uniquely differentiated the two-word adjective-noun phrase condition from the remaining three. This was done by specifying clusters be formed only from the smallest magnitude of three t-values, from the comparisons of two-word phrases vs. two-word lists, two-word phrases vs. one-word phrases, and two-word phrases vs. one-word lists. All three t-tests were performed at each source and time point and the smallest of the three absolute values assigned to each point. Clusters were then formed from these smallest, absolute-valued statistics using the TFCE method, and ten-thousand permutations of condition labels were used to estimate cluster p-values. Notably, due to the use of the absolute values in the first stage of the test, the resulting clusters could contain patches of cortex that showed both negative- and positive-polarity dissociations between conditions, causing the difference within the cluster, when averaged over sources, to be approximately zero. If this was found to be the case, only the positive-polarity component of the cluster was visualized in the results.

Experiment 1 and 2 analyses were performed in lateral surface ROIs. A left anterior temporal lobe ROI was defined as the anterior half of the superior, middle, and inferior temporal gyri, as well as the temporal pole. This ROI was designed to encompass those regions in which previous minimal composition MEG paradigms have identified the effects of interest (e.g., Bemis & Pylkkänen, 2011; Westerlund & Pylkkänen, 2014; Flick et al., 2018). A left posterior temporal lobe ROI was defined as the posterior half of the superior and middle temporal gyri and anterior one third of Brodmann Area 39. This ROI was designed to contain the pMTG site examined by Teige et al., (2019) and implicated in semantic control manipulations by the meta-analysis of Noonan et al., (2013). A left angular gyrus ROI was defined was as the posterior two-thirds of Brodmann Area 39 and the whole of Brodmann Area 40, so as to capture the “automatic semantic processing site” identified in the meta-analysis of Humphreys and Lambon Ralph (2015).

Tests were performed in all three of these ROIs and in three windows defined relative to the onset of the phrasal head (or the second item in list and one-word trials): 150-300 ms, 300-450 ms, and 450-600 ms. These temporal windows were motivated by the previous timing of left ATL composition responses (150-300 ms and 450-600 ms in Bemis & Pylkkänen, 2011, and Flick et al., 2018, respectively) and the inclusion of the intervening time (300-450 ms). We also performed cluster-based permutation tests in the same windows on the mean responses within a functional ROI defined from the letter-string response localizer.

Spatiotemporal cluster-based permutation tests were also used to analyse data collected in Experiment 2 (noun-noun combinations). Tests were performed in the same set of ROIs used in the analysis of Experiment 1 data. Clusters were formed from t-values comparing response to the head nouns in Material and Contents compounds and post-hoc comparisons to the adjective-noun phrases were used to aid the interpretation of the results. All tests were performed in the same time windows used in the analysis of Experiment 1. We note that the presentation of the isolated words in Experiment 2 could be used to create statistical contrasts of responses to nouns in isolation and as phrasal heads (e.g., cabinet vs. trophy cabinet). However, the absence of a placeholder stimulus for the modifier (as in Experiment 1: xkq boat) effectively precludes this possibility, as the comparison would be confounded by differences in the baseline windows. For this reason, we abstained from analyzing the two-vs. one-word contrast in Experiment 2, relying instead on inferences from the appropriately designed contrasted in Experiment 1.

Unless otherwise indicated, p-values in the primary tests of each dataset were corrected for multiple comparisons across ROIs and time windows using the False Discovery Rate (FDR) procedure (Benjamini & Hochberg, 1995), with a corrected significance threshold of p < 0.05. In the letter-string response localizer, this included the tests for Type I and II Noise Level responses, and the String Type response. In the analysis of Experiments 1, this included the spatiotemporal tests in the left ATL, PTL, and AG, as well as the temporal tests performed in the functional ROIs from the letter-string response localizer. In Experiment 2, this included the left ATL, PTL, and AG ROIs, as well as the String Type ventral ATL fROI. Following the reasoning that each observed cluster was representative of its encompassing ROI and time window (see discussion in Sassenhagen & Draschkow, 2019), the smallest cluster p-value in each test window/ROI was submit to the correction procedure to determine if there was sufficient evidence to support a deviation from the null hypothesis.

### Spatial Resolution and Source Crosstalk

The minimum norm approach to the MEG inverse problem is a linear method that can be examined and characterized based on how well it distinguishes activity at individual sources or, conversely, blurs this activity together (Hauk et al., 2019). This blurring is often referred to as “signal leakage”, a problem that is inherent in the minimum norm method. One consequence of this is that the localization of a particular response pattern to a particular region may not be veridical, because estimated responses at that location could be conflated with activity from other locations. One way to examine this leakage is through the crosstalk functions (CTFs) for constituent sources. For each individual, the source estimation procedure requires specification of a forward matrix (based on anatomy and specifying the transformation from source-to sensor-space) and an estimated inverse matrix (specifying the transformation from sensor-to-source-space). CTFs can be extracted from the product of these two matrices, referred to as a resolution matrix (see Hauk et al., 2019 an approachable introduction to spatial resolution metrics, including the derivation of the resolution matrix, for more detailed discussions see Liu et al., 2002; Molins et al., 2008; Hauk et al., 2011). For a particular source *i*, the CTF can be found in the *i*-th row of the resolution matrix, and captures how unit magnitude at each source would leak into the estimate for source *i*. Conversely, if one were interested in how activity at source *i* leaks or spreads into activity at other sources, one could examine the *i*-th column of the resolution matrix, referred to as the point-spread function (PSF) for source *i*.

Here we examined the spatial accuracy and overlap of crosstalk functions for each of the regions identified in the analyses below. For each area, we computed the crosstalk functions for all sources contained within it to determine if the pattern may be generated or influenced by activity elsewhere in the cortical surface. This was particularly important when examining patches of cortex that showed similarly timed effects, as they could be either independent responses or influenced by shared signal leakage. We also examined each region’s peak localization error (PLE). For an individual source, PLE was defined as the distance between the location of the source and the absolute maximum of that source’s point spread function. In an ideal scenario, a source’s PLE is equal to zero (i.e., the location of the maximum is the source itself). For a patch of cortex, the PLE and CTF were computed as the mean over all constituent sources.

## RESULTS

### Behavioral Results

Accuracy data collected in Experiments 1 and 2 were examined for the purpose of confirming attendance to the experimental procedures. Reaction time data were not analyzed as participants would occasionally take breaks before responding to the images if they were feeling fatigued at that point in the experiment. Across both experiments, accuracy was high, indicating that participants understood and attended to the tasks. In Experiment 1, mean accuracy was 95.7% correct (sd = 4.72%). All participants scored above 84% accuracy and only four below 90%. Coding of accuracy in Experiment 2 was more ambiguous due to the increased complexity of the words’ depictions (i.e., steel, titanium canvas, etc.) and the corresponding increases in the complexity and ambiguity of the image stimuli. Nevertheless, mean accuracy was high again (m = 86.04%, sd = 3.05%), with all participants scoring above 78%. No behavioral data were collected in the letter-string response localizer, as participants passively viewed the stimuli.

### Letter-string response localizer

Analysis of data collected in the letter-string response localizer replicated the patterns found in by Gwilliams et al. (2016). For each analysis, multiple statistically significant spatiotemporal clusters were found, however we report here those that matched, most closely, the patterns found in the previous work. The full set of clusters are reported in the Supplementary Materials. Note, this does not mean that the pattern identified in a spatiotemporal cluster was only found at that location (or timing). Indeed, this is a weakness of spatiotemporal cluster tests: one cannot make inferences regarding the location of a cluster within an encompassing ROI nor its timing within an encompassing test window (Sassenhagen & Draschkow, 2019). This is in addition to the inherent uncertainty regarding spatial accuracy of MEG source estimation methods (see below). We do, however, consider clusters that localized to the same regions and showed the same waveform morphology as found by Gwilliams et al. to provide compelling evidence that their specific timing and/or localization are generalizable.

Figure 2 displays the localizer results. For each analysis, cluster-based permutation tests were performed on the time course of regression coefficients. For ease of interpretation, Figure 2 shows the mean response within each relevant condition, computed as the average within the highlighted clusters. In lateral occipital and posterior temporal areas, we observed an initial sensitivity to visual noise properties of the stimuli before any letter vs. symbol string discrimination. A significant effect matching the profile of the original Type I Noise Level effect was found in the lateral and ventral occipital lobe, spanning 100-180 ms after stimulus onset (corrected p = 0.0004), and capturing a negative-going component in response to high noise stimuli that was absent in low noise stimuli. A Type II Noise Level effect, an opposing polarity dissociation between high and low noise conditions, was found in the posterior fusiform gyrus between 100 and 160 ms (corrected p = 0.0004), with the peak dissociation at approximately 130 ms.

Moving more anteriorly and later in time, we observed a change in sensitivity to a discrimination of letters and symbol strings. A significant String Type cluster was found in the anterior fusiform and inferior temporal gyri between 205 and 300 ms (corrected p = 0.0014), showing greater positive responses to letters relative to symbols. Inspection of the cluster’s waveforms (Figure 2, middle-right) revealed an earlier peak at approximately 170 ms after stimulus, which showed numerically greater responses to letter-strings than symbols, although this did not reach statistical significance in post-hoc tests.

Expanding on the analyses performed by Gwilliams et al., and repeated here, we also performed a series of follow-up tests that separated the four- and one-unit letter and symbol-string stimuli. String Type regression analyses using only four-unit trials (4-letter words vs. 4 symbols) and only one-unit trials (1 letter vs 1 symbol) both yielded statistically significant effects (1-unit: 240-300 ms, uncorrected p = 0.0009; 4-unit: 205-300 ms, uncorrected p = 0.0008). Notably, the largest cluster identified in the contrast of 4-unit stimuli was located more anteriorly than that found in the contrast of 1-unit stimuli, as well as the original cluster that was found when both lengths were included in the analysis. Supplementary Figure 1 displays the overlap of these clusters on the ventral surface. Although the precise locations of the clusters were not subject to a statistical test and therefore should not be over-interpreted, we examined this pattern further by performing a linear regression analysis. A linear model was fit to assess the relationship between posterior-to-anterior position along the left fusiform gyrus and the mean difference between responses to letter and symbol stimuli between 200 and 300 ms after stimulus onset. This revealed a significant positive relationship (b = 3.22, Intercept = 0.12, R^2^ = 0.49, p < 0.001) for only the 4-unit stimuli, with words elicited increasingly more positive responses than their symbol-string counterparts as one moved anteriorly along the fusiform gyrus. The same analysis of single-unit stimuli (letters vs. single symbols) was not significant (b = 0.698, Intercept = 0.187, p > 0.69).

Finally, we also asked whether the largest significant cluster identified in each separate String Type test distinguished between the letter and symbol stimuli of the other length. This was done by conducting a temporal cluster-based permutation test on the mean estimated source amplitude within each cluster. The results of these tests are in Supplementary Figure 1. Although the more posterior cluster identified in the 1-unit contrast (1 letter vs. 1 symbol) did not show a significant difference between 4-unit stimuli (word vs. 4 symbols) in the original analysis window (130-300 ms), a clear dissociation did appear later the epoch (100-600 ms test window, significant clusters: 300-405 ms, uncorrected p = 0.0076; 415-450 ms, uncorrected p = 0.0043). On the other hand, the more anterior ROI that distinguished 4-unit words and symbol strings, at 205-300 ms, showed no differences between the 1-unit letters and symbols at any point in the epoch.

The spatial extents of the clusters that matched the Gwilliams et al. (2016) results were adopted as group-level functional (f)ROIs in the analyses reported below to examine where along this series of responses we might observe a sensitivity to the different word combination conditions. We report the results using only the general String Type fROI, defined across letters- and four-letter words compared to length-matched symbol stimuli, rather than the overlapping but more anterior cluster found in the contrast of only words and four-unit symbol strings. The same general pattern of results was found if the more anterior word vs. 4-symbol cluster was used instead.

### Experiment 1: Adjective-noun combinations

Spatiotemporal cluster-based permutation tests were first performed in the three ROIs defined as the left anterior temporal lobe (ATL), posterior temporal lobe (PTL) and angular gyrus (AG). Statistically significant composition effects (i.e., a dissociation of the two-word phrase condition from the remaining three) were found in all three ROIs. In the left ATL, significant clusters appeared in the earliest and latest time windows (150-300 ms, p = 0 .0063; 450-600 ms, p = 0.171; all reported p-values FDR-corrected) while in the left PTL ROI significant clusters were found in all time windows (150-300 ms, p = 0.0006; 300-450 ms, p = 0.0081; 450-600 ms, p = 0.0006). In the left AG ROI, significant clusters were found in the 300-450 ms (p = 0.0081) and 450-600 ms (p = 0.0088) windows. Figure 3 shows the time-course of activity localized to representative clusters in each ROI. In the left PTL, the significant cluster in the early time window contained nearly symmetrical negative and positive polarity dissociations amongst the conditions (i.e., mirrored about zero), which caused the region’s time-course to show no dissociation when responses were averaged across all sources. Thus, for visualization in Figure 3, the cluster was masked using those sources that showed a net positive difference between two-word phrases and two-word lists. In the ATL and PTL, dissociations between the two-word phrases and the remaining conditions were notably similar and the largest significant cluster in the left ATL ROI (shown in Figure 3) was located along the superior temporal gyrus (STG), just adjacent to the left PTL cluster. This is discussed further below.

**Figure 3.**
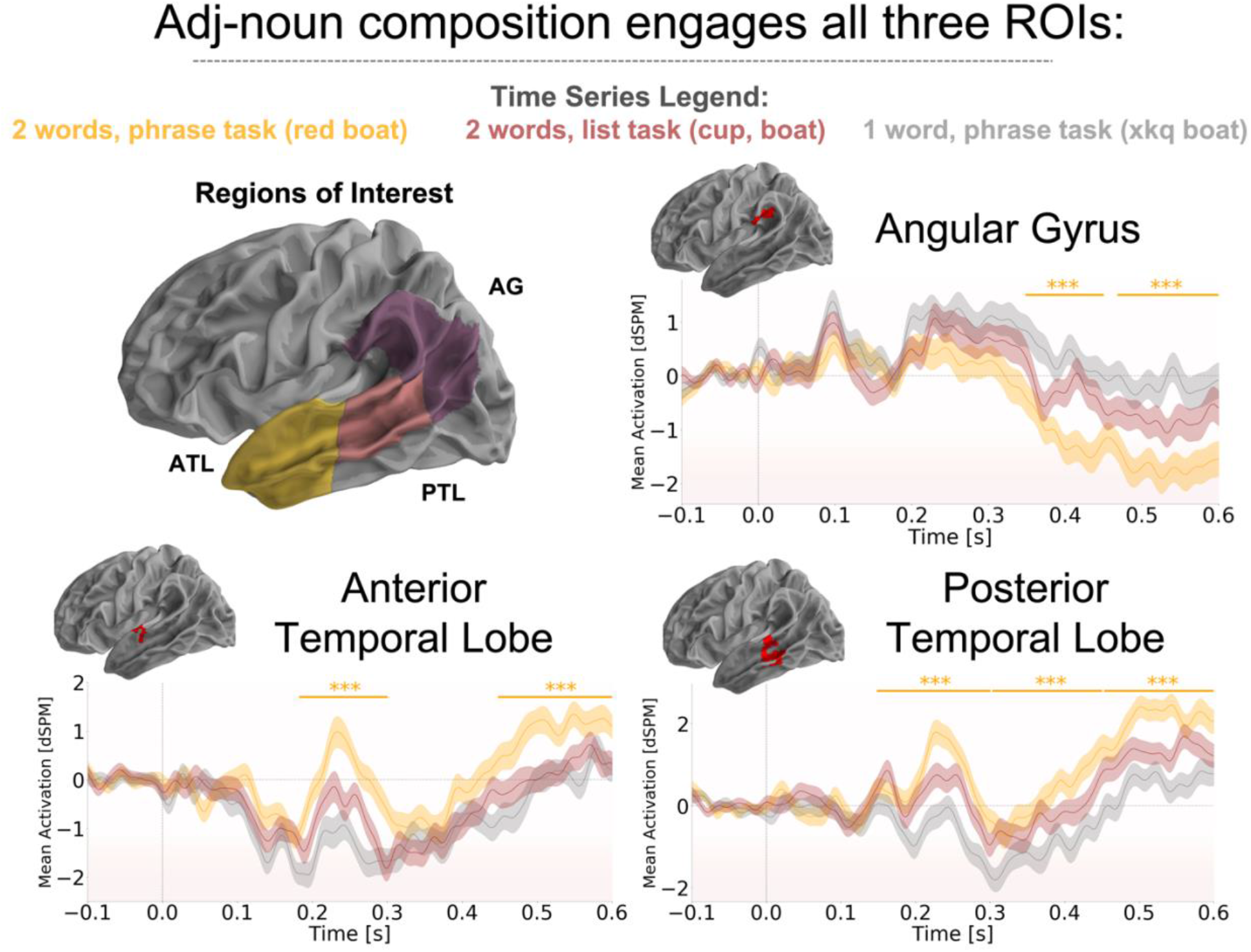
Experiment 1. Adjective-noun composition. Responses were examined in three regions of interest (ROIs, top-left): the left anterior temporal lobe (ATL), posterior temporal lobe (PTL), and a temporal-parietal region encompassing the angular gyrus (AG). Significant composition effects, defined as magnitude increases in response to two-word phrase (e.g., red boat) relative to two-word lists (cup, boat) and one-word baseline materials (xkq boat), were found in all three ROIs. In each time series, 0 ms indicates the onset of the second word on each trial (i.e., the phrasal head). Shown here are representative clusters from each region. Stars and lines indicate significant composition effects. In the ATL (bottom-left), significant effects were found in both the 150-300 ms and 450-600 ms windows, while the AG (top-right) showed the effects in the 300-450 ms, and 450-600 ms windows. The left PTL (bottom-right) showed composition effects in all three analysis time windows. Although the one-word list condition is not shown here for ease of visualization, all composition effects, by test definition, also showed a dissociation from this fourth condition.

We next examined responses in the functional ROIs defined from the letter-string response localizer, using temporal cluster-based permutation tests to compare mean responses within each region. Of particular interest was where along this posterior-to-anterior progression we may observe the earliest sensitivity to conceptual-combination. Figure 4 displays the results. In the anterior fusiform ROI that differentiated letter-strings and symbols, statistically significant composition effects were found in all three analysis windows (150-300 ms, p = 0.0050; 300-450 ms, p = 0.0023; 450-600 ms, p = 0.0006). In contrast, the Type I and II Noise Level regions showed markedly consistent patterns across all conditions, with no statistically significant differences across conditions (all p-values > 0.0780).

**Figure 4.**
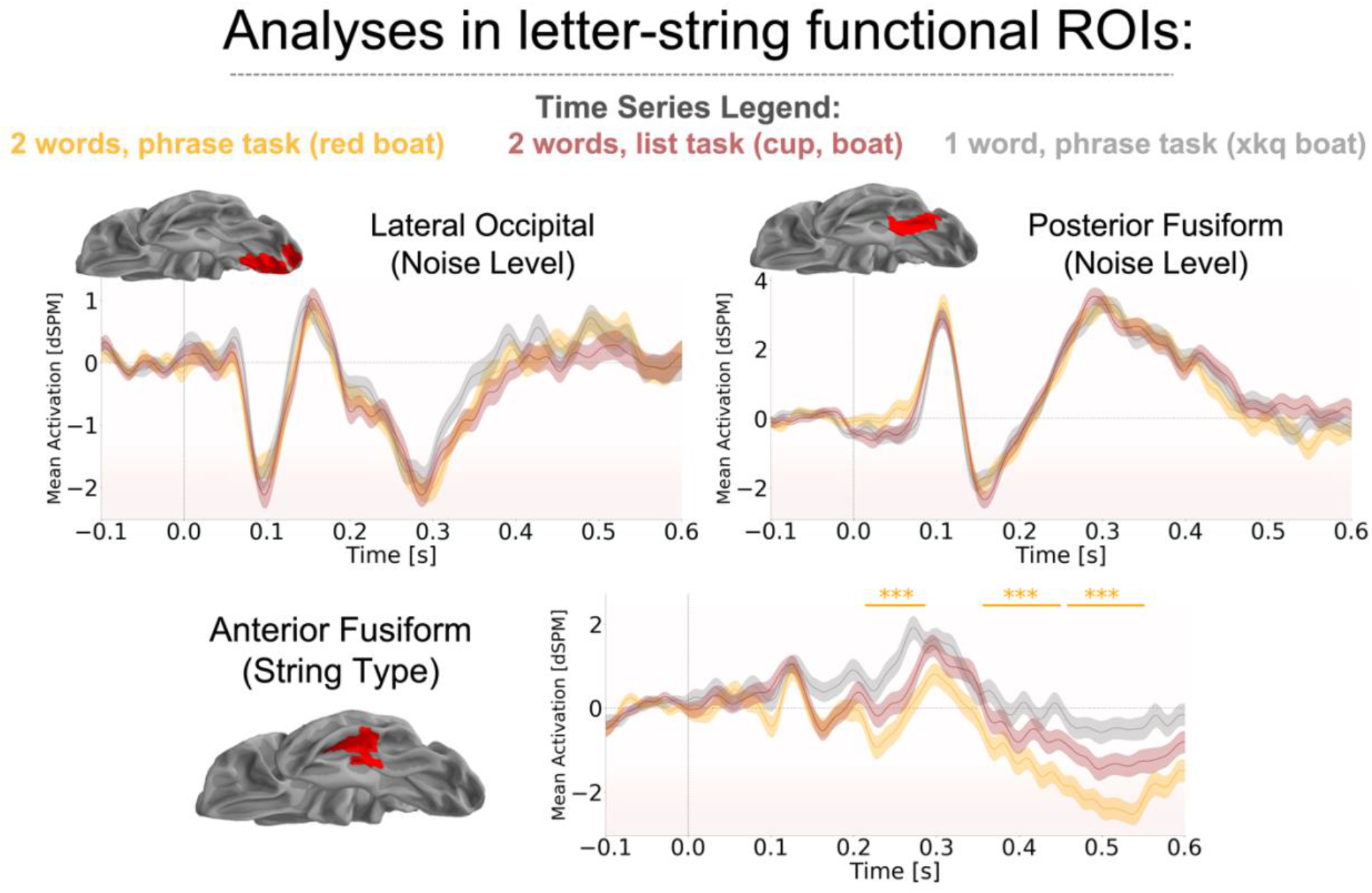
Experiment 1. Analysis of functional ROI responses defined from the letter-string response localizer. In the lateral occipital and posterior fusiform ROIs, shown to be sensitive to the level of visual noise in letter- and symbol-string stimuli, all conditions elicited markedly consistent responses. In the anterior fusiform ROI, composition effects were found in all three analysis windows. In each time series, 0 ms indicates the onset of the second word on each trial (i.e. the phrasal head). Stars and lines indicate significant effects.

In summary, the results of Experiment 1 demonstrated that ROIs capturing the left anterior temporal cortex (150-300 ms, 450-600 ms), posterior temporal cortex (150-600 ms), and angular gyrus (300-600 ms), all contained estimated source amplitudes implying a contribution to adjective-noun composition. Additionally, a left anterior fusiform and inferior temporal ROI that was defined based on a discrimination of letter-strings and symbols (between 130-300 ms) also showed increased response magnitudes to two-word phrases relative to the list and single word conditions, beginning in the 150-300 ms window following the phrasal head. This was the only ROI defined from the letter-string response localizer that showed a sensitivity to phrasal composition.

### Experiment 2: Noun-noun combinations

The results of Experiment 2 are shown in Figure 5. Spatiotemporal cluster-based permutation tests were performed in all three primary ROIs to identify responses modulated by the two types of noun-noun compounds: Contents compounds, involving a spatial or functional thematic relation between the composing words (e.g., trophy cabinet), and Material compounds, where the modifier specified the material that the head noun was made of (e.g., metal cabinet). The only region that showed a significant modulation of responses by Compound Type was the posterior temporal lobe ROI, in the 150-300 ms window (largest cluster: 258-298 ms, p = 0.0058). This ROI also contained multiple clusters in the later test windows, however these did not reach the threshold for statistical significance after multiple comparison correction. These tended to overlap, spatially, with the 150-300 ms window cluster, capturing the prolonged dissociation apparent in the waveform in Figure 5. Estimated activity in the significant cluster showed a pronounced positive peak in response to head nouns combined in a spatial/functional relationship with another noun (e.g., trophy cabinet), which was of greater magnitude than responses to nouns modified by a material (e.g., metal cabinet). Follow-up t-tests of mean responses within the cluster confirmed that Contents Compounds also elicited greater magnitudes than the color-noun phrases (t(27) = 3.52, p = 0.002), with no significant difference found between Material Compounds and color-noun phrases (t(27) = −1.71, p = 0.098).

**Figure 5.**
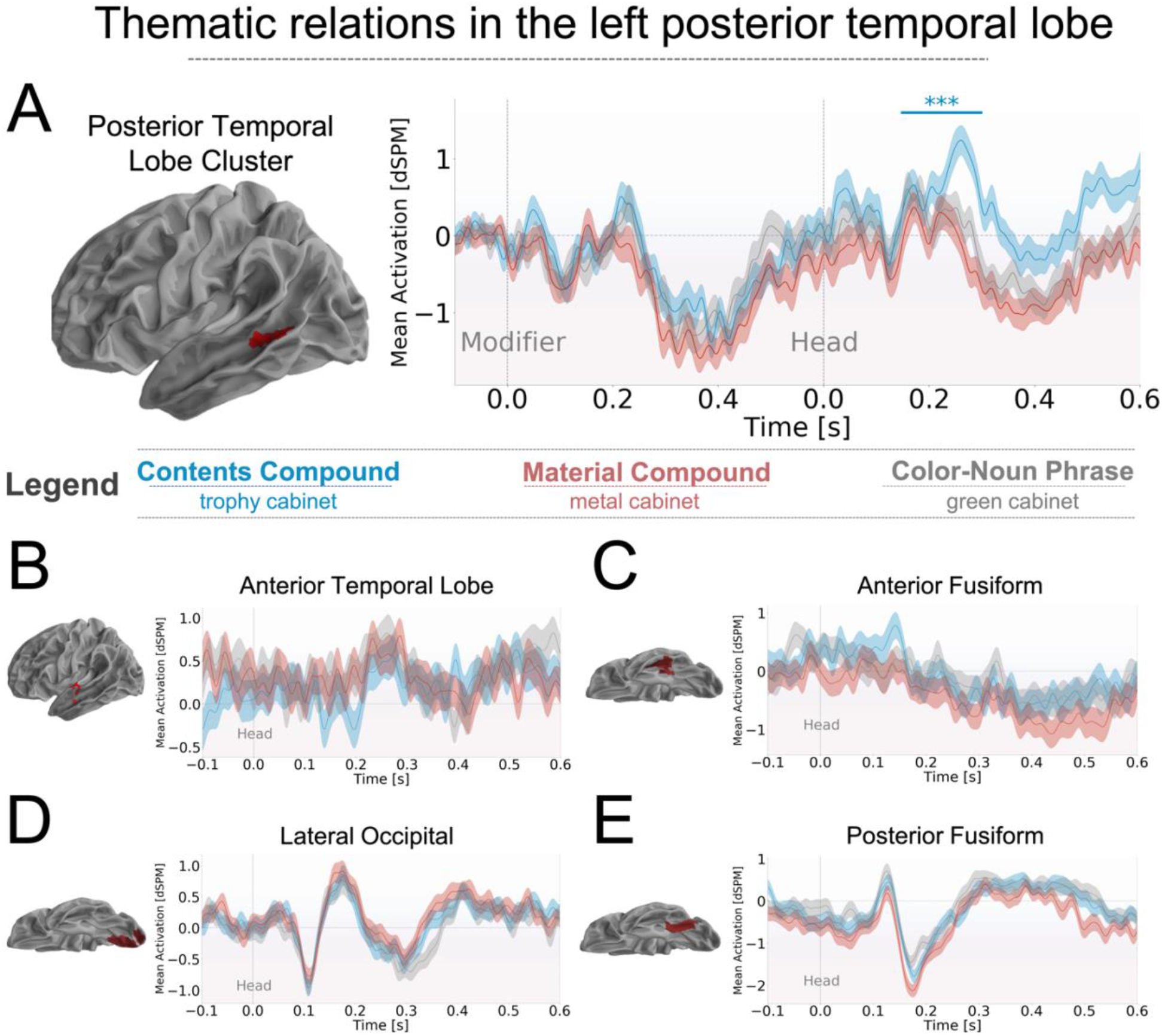
**Experiment 2.**(A) Contents compounds, involving a spatial or functional relation between the combined nouns, elicited increased responses in the left posterior temporal ROI, in the 150-300 ms test window. This increase was relative to both the Material Compounds and color-noun phrases. (B) Cluster tests in the left ATL ROI failed to find any significant difference in responses to three types of stimuli. Shown here is the mean response in the adjective-noun cluster identified in Experiment 1. Tests in the Angular Gyrus ROI also failed to find significant differences. (C) - (E) None of the ROIs defined from the letter-string response localizer showed a significant difference between the two Compound Types

No significant differences were found between conditions in any of the functional ROIs defined from the letter-string response localizer, including the anterior fusiform ROI that showed a main effect of adjective-noun composition in Experiment 1. In follow-up analyses, we treated the adjective-noun composition clusters found in the left ATL, PTL and AG ROIs in Experiment 1 as functional ROIs and performed cluster-based tests of responses averaged over the sources contained in them. This uses the adjective-noun contrast of Experiment 1 as a group-level functional localizer, identifying sites of sensitivity within each encompassing ROI. Once again, no significant differences between the Experiment 2 conditions were found in these analyses. Although not identifying any differences between the two compound types, exploratory tests contrasting the Material compounds and color-noun phrases in the two Noise Level ROIs identified one (uncorrected) significant cluster in each. This was found in the 300-450 ms window (cluster: 327-368 ms, uncorrected p < 0.0034) in the Noise Level I ROI, and the 150-300 ms test window (cluster: 169=192 ms, uncorrected p < 0.0143) in the Noise Level II ROI. Since these represent secondary findings, for which we had no a priori theoretical predictions, we do not discuss them further.

### Localization Accuracy and Crosstalk

To examine localization accuracy and the blurring of sources across the loci of our effects, we computed the average peak localization error (PLE) from the point spread functions of sources within each of the spatiotemporal clusters identified in the previous analyses. This was done using the covariance matrices and inverse estimators for all participants from data collected in Experiment 2. The results are shown in Figure 6, with the removal of one outlier participant (mean error = 2.28 cm across ROIs) for visualization. Overall, the localization error was within an acceptable range, with the mean values across participants falling between 0.88 cm and 1.60 cm, and with no one region showing a markedly greater localization error than the others. With the exception of the posterior fusiform, the general pattern was that more anterior temporal ROIs tended to show greater localization errors, consistent with previous results finding particularly high localization uncertainty in the anterior temporal lobe (Hillebrand & Barnes, 2002; Hauk et al., 2011). The mean PLE was lowest in the lateral occipital region ROI (m = 0.88 cm, sd = 0.15 cm), followed by the posterior middle temporal gyrus cluster that showed a sensitivity to Compound Type (m = 0.98 cm, sd = 0.23 cm), then two clusters that showed adjective-noun composition effects in Experiment 1: the angular gyrus (m =1.05 cm, sd = 0.21 cm) and anterior superior temporal gyrus (m = 1.39 cm, sd = 0.40 cm). Finally, the locations with the greatest localization error were the posterior (m = 1.47 cm, sd = 0.39) and anterior (m 1.60 cm, sd = 0.49 cm) fusiform ROIs defined from the letter-string response localizer.

**Figure 6.**
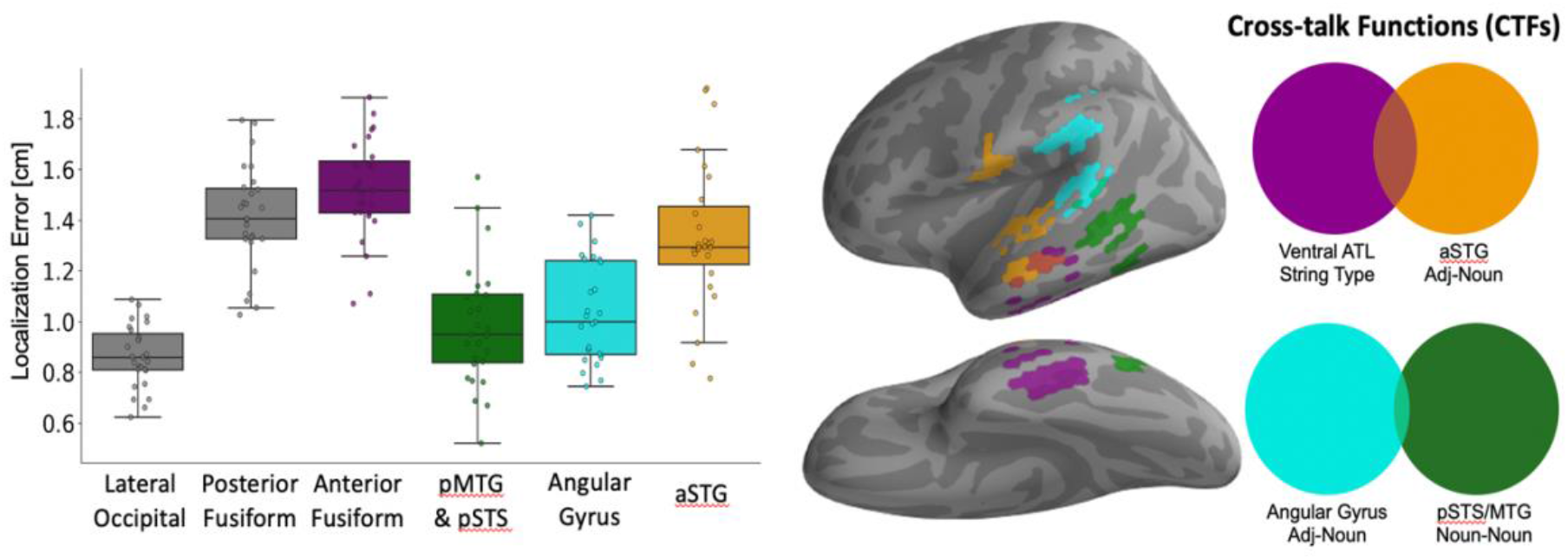
Left: Peak localization error (PLE) was calculated as the Euclidean distance between the position of each constituent source and the absolute maximum in that source’s point spread function. PLE for an ROI was calculated as the average PLE of all constituent sources. With the exception of the posterior fusiform ROI, PLE tended to increase with anteriority. Values shown here are after the removal of one outlier participant. Right: An examination of the mean crosstalk functions from the anterior fusiform and anterior superior temporal gyri revealed notable overlap between the regions’ CTFs. The mean CTF for the AG and pSTS/MTG regions provided no evidence that responses here were conflated with source activity falling in the other two ROIs, and the two regions showed little overlap. String Type indicates clusters that differentiated letter-strings from symbol-strings; Adj-Noun indicates clusters implicated in adjective-noun combination; Noun-Noun indicates clusters implicated in noun-noun combination. ATL = anterior temporal lobe; aSTG = anterior superior temporal gyrus; pMTG = posterior middle temporal gyrus; pSTS = posterior superior temporal sulcus.

In more targeted examinations we first compared the crosstalk functions for all sources in four of the identified clusters: (i) the ventral anterior fusiform region that showed the effect of String Type, the anterior superior temporal gyrus (aSTG) and angular gyrus clusters that showed effects of adjective-noun composition (ii & iii), and (iv), the posterior superior temporal sulcus/gyrus (pMTS/MTG) region that distinguished the two noun-noun compound types. Of particular interest was whether responses localized to each of these regions were contaminated by leakage from the others, especially when two areas showed similar response patterns. The CTFs for constituent sources in each ROI were averaged, converted to absolute values, and masked with a half maximum threshold. As shown in Figure 6 (right), ventral ATL and left aSTG clusters showed notable overlap among their CTFs in the lateral temporal lobe. This appears to explain the apparent mirroring of the anterior fusiform and STG responses in Experiment 1, which showed similar patterns with opposite polarities (compare Figures 3 & 4). Moreover, it suggests that caution is needed when considering whether MEG-localized responses in the ventral and superior ATL capture separable and independent computations in the comprehension of adjective-noun combinations. The shared response pattern appears, at least in part, to be an artifact of the source estimation method. Overlap was also found between the adjective-noun cluster in the angular gyrus and left anterior temporal lobe, in posterior sections of the STG. The CTFs of the AG and noun-noun composition pMTG clusters, on the other hand, were largely disjoint from each other, and non-overlapping with the two anterior regions.

Second, we also assessed the crosstalk overlap for those clusters identified in the temporo-occipital progression of visual noise and letter-string responses. CTFs were extracted from the lateral occipital noise level cluster, posterior temporal noise level cluster, and the anterior fusiform string type cluster, found from the contrast of words and symbols strings. These are displayed in Supplementary Figure 2. The masked CTFs of these clusters showed only minimal overlap, found between the lateral occipital and posterior fusiform responses. The anterior string type cluster’s CTF did not overlap with either of those of the more posterior regions. While variability in CTFs from dataset to dataset is unclear at this time, this finding provides at least one datapoint suggesting that magnetoencephalography recordings and linear source estimation can spatially discriminate (at the group level) word-level and visual/orthographic processing in left occipital and ventral temporal cortex.

### Whole-brain statistical contrasts

Finally, to ensure that a focus on ROI analyses did not lead to missed dissociations in other cortical regions, we examined statistical contrast maps between the primary conditions of Experiment 1 and 2 across the cortical surfaces. We note that these are primarily descriptive results, not subject to appropriately corrected statistical tests for their robustness. The lateral and ventral surface maps are shown in Supplementary Figures 3-5. In the comparison of two-word phrases and two-word lists from Experiment 1, in addition to the left ATL, AG, and PTL dissociations found in the ROI analyses, dissociations were also seen in the right anterior temporal lobe (400-600 ms), prefrontal cortex (200-300 ms), and temporal-parietal cortex (500-600 ms). The whole-cortex visualizations also make clear the prominence of the ventral ATL dissociation between two-word phrases and two-word lists (as well as the one-word items), which can be seen as a sustained cluster of t-values, particularly from 400 ms onwards. In the comparison of Contents and Material Compound stimuli of Experiment 2, additional differences between the conditions were found in the mid fusiform gyrus, toward the medial surface (100-400 ms), the right anterior temporal pole (100-200 ms), and dorsal sections of the right parietal lobe (200-300 ms & 500-600 ms). On the medial surface, a dissociation between the two compound types was also seen in the posterior cingulate gyrus, bilaterally, as well as the isthmus of the cingulate gyrus (see Supplementary Figure 4). The comparison of Contents Compounds and adjective-noun phrases (i.e., trophy cabinet vs. green cabinet) revealed notable differences in the vicinity of the left inferior frontal gyrus (200-300 ms), the medial fusiform gyrus (200-300 ms), and the right anterior temporal lobe (100-300 ms).

## DISCUSSION

This work examined how the comprehension of particular types of adjective-noun and noun-noun concepts differentially taxed neural responses in visual word processing and candidate semantic hub regions. Figure 7 displays a summary of the primary findings. Our results demonstrate that posterior sections of the left temporal lobe, encompassing the posterior middle temporal gyrus, superior temporal gyrus, and superior temporal sulcus, responded with greater magnitudes to the head of a noun-noun combination when that combination required a thematic relation between the composing words (e.g., trophy cabinet is a cabinet that holds trophies). This finding follows previous work implicating the left posterior temporal lobe in the processing of verb and event knowledge (e.g., Bedny et al., 2008, 2014). Since this knowledge is related to how entities co-occur in events, or as verb arguments, and this also makes up one’s thematic knowledge of entities (i.e., how they occur or complement one another in various scenarios), the pattern suggests a shared computation housed in the left posterior temporal lobe. We also found, however, that this left PTL region was recruited in the comprehension of adjective-noun phrases when the adjective modified only the color of the head noun (e.g., red boat), demonstrating that its contribution to composition is not limited to relational processing. Both this effect, and the thematic relation effect, were found in the same window relative to the onset of the head noun (150-300 ms). Sections of the left anterior temporal lobe (150-300 ms) and a temporo-parietal ROI containing the left angular gyrus (300-450 ms), were also engaged by color-noun composition.

**Figure 7.**
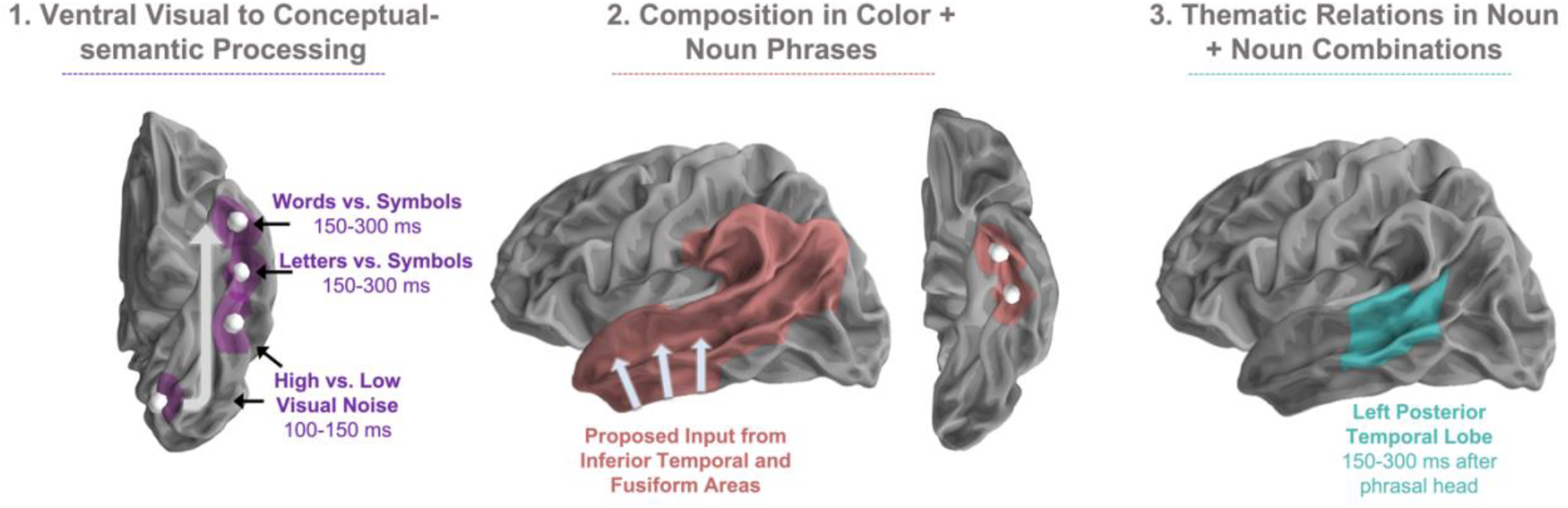
Summary of the primary findings. (1) The results of the letter-string response localizer suggest a visual to conceptual-semantic transformation along the ventral surface of the left hemisphere in support of visual word recognition. White circles indicate the center of mass of each relevant cluster, which showed similar localizations and morphology to those identified by Gwilliams et al., (2016). Shaded purple regions indicate a spherical region with a radius of 15 mm grown around each center of mass. The grey arrow indicates the inferred flow of information processing, from posterior to anterior. (2) Conceptual combination in color + noun phrases elicited the engagement of a distributed left Perisylvian network, along with anterior sections of the left fusiform gyrus. All sections of this network, with the exception of the left temporoparietal ROI containing the angular gyrus, appeared to come online by 150-300 ms after the onset of the phrasal head in the combination. On the basis of the timing of responses along the ventral surface, we propose that this lateral network receives input from inferior temporal and fusiform areas responsible for the visual to conceptual-semantic processing. This is depicted by the grey arrows. (3) Of those areas engaged by color + noun combinations, only the left posterior temporal lobe ROI was further modulated by the need to retrieve/specify thematic relations between composing words in noun-noun combinations. This also appeared between 150-300 ms after the phrasal head. This suggests that the left PTL region houses a functional sub-component related to the retrieval/processing of thematic knowledge, within the larger left hemisphere network that is engaged by relatively simple (e.g., color + noun) word combinations.

Altogether, this pattern suggests that a distributed left perisylvian network comes online to support composition, even when only two words are combined, while a sub-component of this network, the left posterior temporal lobe, also houses computations specifically related to retrieving or otherwise processing thematic knowledge relevant to the ongoing combination of words. Our data also suggest that sections of the left anterior fusiform and inferior temporal gyri (or in their vicinity, see below), are among the first areas along the ventral visual word processing stream to become engaged by composition, which may then feed into this distributed lateral network. Specifically, we observed that a left anterior fusiform area contributed to both visual word recognition (greater response magnitudes to letters than symbols) and adjective-noun combination (greater response magnitudes to two-word phrases than lists or single words), beginning between 150 and 300 ms after word onset. We outline each of these findings in greater detail below, alongside proposed interpretations that we hope will motivate future research.

### Relational processing modulates responses in the left posterior temporal lobe

Our characterization of the posterior temporal lobe’s contribution to noun-noun compound comprehension stems from the thematic and non-predicating nature of the Contents Compounds (e.g., trophy cabinet) as compared to the Material Compounds (e.g., steel cabinet) and adjective-noun phrases (e.g., green cabinet). Specifically, the spatial or functional relationship between the two nouns in Contents Compounds can be plausibly said to rely on thematic knowledge of the concepts (i.e., co-occurrence or complementary roles of the two objects in an event/schema; Estes, Golonka, & Jones 2011) and cannot be paraphrased as a simple predication (i.e., a trophy cabinet is not a cabinet that is a trophy). On the other hand, both Material modifiers and the color modifiers in the adjective-noun phrases can be adequately paraphrased as predications and specify simple (visual) features of the head nouns’ denotations. Heightened response magnitudes to Contents Compounds relative to Material Compounds were observed despite the stimuli being balanced on overall bigram frequency, transition probability from the modifier to the noun, and subjective familiarity as measured by a stimulus norming study.

Considered from this perspective, the present result is consistent with some predictions of dual hub accounts of semantic knowledge (de Zubicaray et al., 2013; Schwartz et al., 2011), as the presumed presence of thematic linkages between words only modulated responses in a posterior region of the left temporal lobe, rather than the left anterior temporal regions proposed, in alternative accounts, to serve both thematic and taxonomic knowledge retrieval. However, we also observed that relative to non-combinatory stimuli, feature modifications in adjective-noun phrases (e.g., red + boat) engaged the posterior temporal lobe and the left angular gyrus ROIs, demonstrating that the contribution of responses in these areas to compositional operations is not exclusive to thematic information processing. Thus, our results do not straightforwardly support the strictest version of a dual hub account, which would posit that feature-based modifications only modulate left ATL responses (although see below for more discussion on separate functions in each region).

The particular localization of the noun-noun compounding effect, in the posterior temporal lobe ROI rather than an ROI containing the angular gyrus, is also noteworthy for its relation to previous findings. The posterior MTG has been found to house heightened responses to verbs relative to nouns (Bedny et al., 2008; Bedny et al., 2014; Bedny & Thompson-Schill, 2006; Davis et al., 2004; Kable et al., 2002, 2005; Martin et al., 1995; Yu et al., 2011, 2012; c.f. Williams et al., 2017) while distributed activations in neighboring sections of the left superior temporal sulcus and gyrus have been found to encode relational roles (e.g., woman as agent, girl as patient) in sentence comprehension (Frankland & Green, 2015, 2020). Assuming that verbs activate event and relation knowledge in memory and considering thematic knowledge as being related to how entities co-occur in events or schemata (Estes et al., 2013), these findings suggest an underlying thematic or relational processing role for the left posterior temporal lobe. Nevertheless, in their comparison of relational (i.e., thematic) and attributive noun-noun compounds, Boylan et al. (2017) found greater hemodynamic response magnitudes for relational compounds in the left and right angular gyri, building on previous work that showed distributed activation patterns in the AG correlated with similarity ratings of verb phrases (Boylan et al., 2015).

Since they are conflicting, these findings suggest, a priori, that the presence of thematic relations in our current stimuli could plausibly be expected to modulate responses in either of the left AG or posterior temporal lobe ROIs. Although we cannot be certain of the precise localization, due to the spatial uncertainty inherent in our source estimation (see Figure 6), our results suggest that it is the posterior MTG, STS, and/or STG regions of the PTL ROI, rather than the AG, that contribute to the processing of thematic relations in familiar noun-noun combinations. This represents a deviation from the AG localization of Boylan et al.’s (2017) relational compounding effect, which was found using fMRI and a different task, and with a set of stimuli that contained many unfamiliar or novel noun-noun combinations. Future work will further inform our understanding of this difference by directly comparing familiar and unfamiliar noun-noun combinations, using a common neuroimaging modality.

The finding that both color + noun and thematic noun + noun combinations engaged the left PTL, relative to their corresponding baseline conditions, raises the question of whether there are shared or distinct computations underlying the region’s involvement in each case. On the one hand, as summarized above, there is now a large body of converging evidence behind the hypothesis that sections of the PTL play a role in relational and/or thematic knowledge processing. However, there is also accumulating evidence for the left PTL’s involvement in syntactic processing in language comprehension (Flick & Pylkkänen, 2020; Matchin, Brodbeck, Hammerly, & Lau, 2018; Matchin, Hammerly, & Lau, 2017; Rodd, Longe, Randall, & Tyler, 2010; Rogalsky et al., 2018; Snijders et al., 2008; Tyler, Cheung, Devereux, & Clarke, 2013). The latter could account for the region’s heightened responses to two-word phrases relative to lists and single words, since the latter two conditions do not require the construction of phrasal structure. It is also intriguing to consider that thematic or relational noun-noun compounds could be place greater demands on lexico-syntactic processing relative to non-relational noun-noun combinations, and this could account for part of their increased activation. This may be the case if, for example, accessing the covert relations between nouns requires specification of an argument structure related to knowledge of an intervening verb (e.g., the cabinet containing the trophies). It should be noted however, that if the PTL is involved in syntactic operations, such as building the structure of a phrase, one would expect it to show effects whenever there is a contrast of phrasal vs. non-phrasal stimuli. Historically, this is not an effect that previous MEG studies of modifier-noun composition have observed, with the finding only appearing in one of the prior studies adopting this paradigm (Bemis & Pylkkänen, 2013). The reasons for this inconsistency are presently unclear, and there remain many intriguing questions regarding how relational and syntactic processing may coexist in the left posterior temporal lobe.

We do note that there is one more pertinent alternative that may account for the PTL’s differential responses to our noun-noun phrases, based on the proposals of the Controlled Semantic Cognition (CSC) framework. As outlined in the Introduction, this account posits that the posterior MTG, along with the left inferior frontal gyrus, contributes to semantic cognition by retrieving contextually relevant but non-dominant aspects of knowledge (Jefferies et al., 2020; Teige et al., 2019). It also specifies that the angular gyrus, in contrast to the posterior MTG, supports more automatic semantic retrievals (e.g., retrieving the dominant rather than subordinate meaning of lexically ambiguous words), while the left ATL functions as the primary semantic hub region.

In light of the Compound Type effect’s location in the PTL rather than the AG, it is worth considering that the difference between the Contents and Material compound conditions, as well as the Contents Compounds and adjective-noun phrases, may be related to semantic control demands rather than (exclusively) thematic vs. non-thematic semantic relations. This idea is even more relevant based on the Experiment 2 whole-brain comparison of Contents Compounds and adjective-noun phrases, which revealed a dissociation between the two conditions in left inferior frontal cortex in the same window as the PTL engagement (though this dissociation was not apparent in the whole-brain comparison with Material Compounds). This area is proposed to also function in semantic control, along with the PTL, in the CSC framework. Notably, although the Contents and Material noun-noun stimuli were balanced on familiarity and frequency properties, the two could differ with regard to semantic control demands if, for example, the typical cabinet one brings to mind when retrieving the word’s meaning is more similar to one’s mental representation of a metal cabinet than one’s mental representation of a trophy cabinet (see relevant behavioral findings from Gray & Smith, 1995). While this seems plausible, at least for these example stimuli, it remains to be tested in future work, which could manipulate both semantic control demands and semantic relations to pit the two explanations against one another.

### Visual word recognition and adjective-noun composition

Although the primary focus of this investigation was the processing of semantic relations between words, our results also suggest implications for how the visual processing of words and letter-strings feeds into conceptual combination operations, particularly color + noun combinations. Consistent with the original findings from the development of the letter-string response localizer (Gwilliams et al., 2016 following Tarkiainen et al., 1999; and see Neophytou et al., 2017), we identified a series of responses along the ventral surface of the left hemisphere that appear to underpin visual letter and word processing. These initially discriminated between high and low visual noise properties of the stimuli, in the lateral occipital and posterior fusiform, and then differentiated letter- and symbol-string stimuli in the middle to anterior fusiform gyrus, with words exclusively diverging from length-matched symbol strings (while single letters did not) in the most anterior sections of the left fusiform. This sequence of responses thus replicates previous evidence (e.g., Gwilliams et al., 2016; Vinckier et al., 2007; Woolnough et al., 2020) for a transition along the ventral surfaces of left occipital and temporal cortex from visual and orthographic to lexical processing.

At the more anterior sites of this sequence, in the mid and anterior fusiform, the timings of the effects were noteworthy for two reasons. First, there were subtle differences from the previous work using this localizer. Specifically, Gwilliams et al., (2016) observed discrimination between letter-strings and symbols at an earlier latency (150-200 ms) than was observed here (onsetting at 205 ms). From inspection of the waveforms of each study, however, it would appear that the component identified by Gwilliams et al. was present in the earlier peak in our anterior fusiform cluster (see Figure 2D & E, at approximately 170 ms), trending in the same direction as the original effect but not reaching the threshold for statistical significance. Second, the extended durations of the anterior fusiform discrimination between letters and symbol strings raises questions about what type of computations would result in divergences this long. Supplementary Figure 1 demonstrates that these latencies appeared to differ between the letter and word contrasts with their length-matched symbol strings, such that words diverged from symbols longer than the letter stimuli. In the most anterior fusiform ROI, the waveforms suggested a divergence of at least 200 ms between words and symbols. In an examination of mid-fusiform responses to words and word-like stimuli, Woolnough et al., (2020) found that, in sentence reading, responses to high frequency words diverged from pseudowords by approximately 180 ms after onset, for a duration of upwards of 300 ms, congruent with the present findings. Low frequency words, on the other hand, did not diverge from pseudowords until later in the epoch, which the authors interpreted as evidence that the mid-fusiform maps visually presented word inputs to entries in the mental lexicon. Our findings are consistent with this proposal, and further implicate the site of this mapping, the left mid and/or anterior fusiform gyrus, in the combination of word meanings.

This proposal stems from our examination of how the letter- and word-processing areas responded to the adjective-noun and noun-noun contrasts of Experiments 1 and 2. Of these, only the left anterior fusiform showed any sensitivity to combinatorial demands. This was exclusively in its responses to adjective-noun stimuli and manifested as relatively early (150-300 ms) dissociations of two-word phrases from two-word lists, and one-word items (see Figure 2). We highlight this as a tentative site for the culmination of a visual-to-conceptual transformation in the anterior fusiform (similar to the proposal of Woolnough et al., 2020), which enables the meaning of a visually presented phrasal head (e.g., boat) to quickly feed into coarse conceptual combination operations (e.g., red + boat). The parallel timing of adjective-noun effects in the posterior and lateral anterior temporal lobe (also appearing between 150 and 300 ms) suggests that related information may be relayed to these perisylvian areas, either from the anterior terminus of this stream or earlier points along it. The finding that responses in this region did not differentiate the two noun-noun compound types (Material vs. Content Compounds) offers that it may represent a relatively early stage in conceptual-semantic processing, able to distinguish only between some compositional and non-compositional stimuli, but not subtleties among them. This is congruent with the results of Ziegler & Pylkkänen (2016), which found that the left ATL’s contribution to phrasal composition, 150-200 ms after the phrasal head, was limited to instances where the modifier did not require in-depth processing of the head noun (i.e., intersective adjectives such as *dead* which have context-independent meanings). Moreover, previous electrophysiological findings have demonstrated that, slightly earlier than the timing of our effects, a feedforward pass of activity along the ventral surface of the left hemisphere supports a transformation from visual word input to coarse semantic category information (Chan et al., 2011). Our results are consistent with this proposal, extending the functional scope of this (presumed) forward pass to include a role in the combination of word meanings.

The anterior fusiform was not, however, the only location at which we observed heightened responses to the conceptual combination of color + noun phrases relative to word lists and one-word items, and our examination of crosstalk functions suggests caution concerning the exact location of this effect. Notably, we also saw the same response pattern in sections of the ATL, PTL, and the ROI containing the left angular gyrus. With the exception of the left AG, all of these effects first appeared in the 150-300 ms window (the AG first showed this dissociation between 300 and 450 ms after the head’s onset). We do note that the left ATL and PTL clusters that passed the threshold for statistical significance were found adjacent to each other, separated only by the border of the two ROIs. On the basis of this, it may be more appropriate to interpret the two as a single mid-superior temporal lobe area that showed sensitivity to adjective-noun composition, accompanying the other sites in the ventral ATL and angular gyrus (though see Sassenhagen & Draschow, 2019 for discussion on the fallacy of interpreting cluster locations). Thus, the present results implicate a more distributed set of left hemisphere regions in the combination of adjective and noun meanings, coming online by 150-300 or 300-450 ms after the onset of a phrasal head. Similar findings have been reported in the hemodynamic literature (e.g., Fedorenko, Hsieh, Nieto-Castañón, Whitfield-Gabrieli, & Kanwisher, 2010; Matchin et al., 2017; Pallier, Devauchelle, & Dehaene, 2011), but to the best of our knowledge this is the first time that the engagement of this larger left hemisphere network has been found in an MEG study adopting the minimal contrast of phrases vs. lists and single words.

### The impact of signal leakage and other limitations

The use of linear source estimation in our MEG analysis comes with important caveats. In our case, examination of the spatial uncertainty in our estimators demonstrated notable blurring or signal leakage across left lateral and ventral temporal lobe areas. This has consequences for the interpretation of responses localized in these areas. In particular, the anterior fusiform ROI, defined from the letter-string response localizer, and the aSTG cluster that showed adjective-noun composition effects (see Figures 3 and 4), overlapped in their crosstalk functions in lateral sections of the temporal lobe surface (Figure 6, right). For each constituent source within a parcel or cluster, crosstalk functions reflect the artificial influence of all other sources on amplitude estimated at that source. Thus, overlapping crosstalk functions suggest that, rather than independent responses, activity localized to each area may reflect a smearing in the transformation of MEG responses from sensor to source space. In other words, instead of two independent locations showing greater response magnitudes to two-word phrases vs. the other stimuli, the ventral ATL and anterior superior temporal gyrus may partially reflect a common pattern present in the sensor-level data, which was artificially smeared across these sections of the temporal lobe in source estimation. As an additional consequence of this smearing, we cannot make strong claims about the precise localization of the sensitivity to adjective-noun composition, which may arise in either the lateral or ventral anterior temporal lobe. In contrast to this, the CTF of the left AG region that was sensitive to adjective-noun composition did not overlap with those of the ventral ATL and aSTG clusters, and the posterior temporal lobe cluster implicated in noun-noun composition showed only minor overlap with the remaining CTFs. This provides more compelling evidence that these localizations are indeed independent, in our source estimators, from responses in the other examined areas.

At a more general level, the present characterization of spatial uncertainty is consistent with previous studies that have found low localization accuracies and dipole detection probabilities in anterior temporal regions (Hauk et al., 2011; Hillebrand & Barnes, 2002; Liu et al., 2002; Molins et al., 2008). Although their dependence on the data mean that CTFs are expected to fluctuate from dataset to dataset, and the degree of variability across studies has not yet been characterized, the overlap of CTFs in the ventral and superior temporal lobe do have specific implications for previous work that has focused on this anterior fusiform region. That is, Gwilliams et al. (2016), using the same functional localizer employed here, found that the activity within this anterior fusiform ROI correlated with the transition probability from stem to suffix in morphologically complex words, suggesting that this region plays a role in morphological decomposition or re-combination (Gwilliams et al., 2016; see also Lewis, Solomyak & Marantz, 2011; Solomyak & Marantz, 2010). The current results highlight that, rather than constrained to the anterior fusiform ROI, one cannot rule out the possibility that this correlation was due to leakage from other cortical areas. Similar concerns can be applied to other previous MEG source analyses focused on this anterior temporal region, including our own work (e.g., Flick et al., 2018). We also note that, although speculative at this time, the heightened spatial uncertainty in this region, and its relation to variability in individual participant anatomy via the specification of a forward model, may account for some of the inconsistencies observed across previous MEG studies of adjective-noun composition, which have identified relevant effects in different sections of the anterior temporal lobe (e.g., Bemis & Pylkkänen, 2011; Flick et al., 2018; Kim & Pylkkänen, 2019; Westerlund & Pylkänen, 2014; Zhang & Pylkkänen, 2015).

Critically, none of this insight would be available without the examination of crosstalk and point-spread functions, reinforcing the notion that these provide a valuable source of information about the quality of linear MEG source estimators, and should be routinely examined (Hauk et al., 2011, 2019). These findings also highlight the caution that should be applied not only in the interpretation of the present results, but also those of future MEG studies that use linear estimators with the goal of identifying functional sub-regions in the anterior temporal lobes.

Another limitation of the present study is related to the number of semantic relations between composing words that were examined in this stimulus set, limited to only “head made of modifier” and “head containing modifier” relations, as well as the color-noun composition. Previous psycholinguistic research (e.g., Gagné & Shoben 1997; more recently, Shmidtke, Gagné, Kuperman, & Spalding, 2018), has examined a much larger number of these semantic relations and pointed out important nuances in them, such as the construal of a constituent concept (i.e., plastic squirrel involves a construal of a squirrel, rather than a live one; Wisniewski, 1996). It will be important for future work to investigate how various relations, requiring different aspects of conceptual knowledge of the constituents, tax neural responses in the left PTL, and/or if these recruit additional cortical areas.

Finally, we note that the present results should not be taken to suggest that only the relatively small number of ROIs examined here underpin all of the computations that enable successful comprehension of multiword concepts or contribute, exclusively, to the precise computations examined (i.e., retrieval of relevant thematic knowledge). Indeed, the uncorrected whole-brain contrast maps suggest the contribution of other cortical areas in each type of composition. In particular, adjective-noun phrases also appeared to dissociate from word lists within the right anterior temporal lobe and prefrontal cortex, two findings that have been occasionally reported in previous studies (Bemis & Pylkkänen, 2011, 2013), as well as sections of the right temporal-parietal junction. Contents Compounds appeared to elicit relatively greater estimated response magnitudes in medial sections of the left ventral temporal lobe (not captured in the letter-string response localizer ROI), and also dissociated from Material Compounds and adjective-noun phrases based on responses localized to the bilateral posterior cingulate and isthmus of the cingulate gyrus (Supplementary Figure 3). Both of these regions have been previously implicated in the processing of noun-noun combinations by Graves et al., (2010), and we leave further refinement of their functional characterizations to future work.

### Conclusions & Future Directions

Understanding a combination of words often requires inserting unstated semantic material between constituents to identify the complete meaning. The present findings suggest that, in reading, when familiar noun-noun combinations involve an implicit thematic relation inserted between them (e.g., a horse barn is a barn where horses are kept), retrieving or otherwise processing this relation leads to the engagement of left posterior temporal lobe areas, by 150-300 ms after the appearance of the phrasal head. This left posterior temporal lobe region, which may be constrained to or extend across the left superior temporal gyrus and sulcus, and left middle temporal gyrus, also appears to be involved in composing the meaning of more straightforward adjective-noun combinations, involving only modification of a concrete noun’s color (e.g., red boat). In this case, our results suggest that the left PTL is recruited alongside superior and ventral sections of left anterior temporal lobe, as well as the left angular gyrus, to support the composition, also beginning by 150-300 ms after the phrasal head. Together, this implies that a distributed left perisylvian network supports these relatively simple word combinations, exemplified by color + noun phrases, while a sub-region of this network, the left posterior temporal lobe, also houses computations specific to the use of thematic knowledge to construct the composed meaning.

Our results additionally position the engagement of these lateral combinatorial areas with respect to visual word processing in occipital and ventral temporal sites of the left hemisphere. Previous and present evidence suggests that, in visual reading, combinatorial operations are first supported by a transformation from orthographic to conceptual-semantic representations that takes place in left lateral occipital, and posterior and anterior fusiform areas. Our data demonstrate that portions of this ventral processing stream show remarkably consistent responses across sequential MEG experiments. They also suggest that anterior fusiform or inferior temporal areas involved in early lexical processing contribute to the combination of word meanings, concurrent with the lateral perisylvian network. We further propose that this early combinatory processing in the anterior fusiform is limited to cases that involve relatively simple, perhaps visual, feature modifications (e.g., red + boat). We hope that future work will further test and refine these proposals by examining word combinations that exemplify a greater variety of semantic relations, and manipulating other theoretically relevant properties, such as semantic control demands and concept typicality, in the context of conceptual combination.

## Acknowledgements

We thank Julien Dirani, Ryan Law, and Benjamin Lang for assistance with MEG data collection. We also thank Haidee Patterson for running the collection of all MRI structural scans, and Brenden Lake and Alec Marantz for comments on earlier versions of this manuscript. This research was partially carried out using the Brain Imaging Core Technology Platform and High Performance Computing resources at New York University Abu Dhabi.

## References

Adachi, Y., Shimogawara, M., Higuchi, M., Haruta, Y., & Ochiai, M. (2001). Reduction of non-periodic environmental magnetic noise in MEG measurement by continuously adjusted least squares method. IEEE Transactions on Applied Superconductivity, 11(1), 669–672. https://doi.org/10.1109/77.919433

Balota, D. A., Yap, M. J., Hutchison, K. A., Cortese, M. J., Kessler, B., Loftis, B., Neely, J.H., Nelson, D.L., Simpson, G.B & Treiman, R. (2007). The English lexicon project. Behavior research methods, 39(3), 445–459. https://doi.org/10.3758/BF03193014

Baron, S. G., Thompson-Schill, S. L., Weber, M., & Osherson, D. (2010). An early stage of conceptual combination: Superimposition of constituent concepts in left anterolateral temporal lobe. Cognitive neuroscience, 1(1), 44–51. https://doi.org/10.1080/17588920903548751

Bedny, M., & Thompson-Schill, S. L. (2006). Neuroanatomically separable effects of imageability and grammatical class during single-word comprehension. Brain and language, 98(2), 127–139. https://doi.org/10.1016/j.bandl.2006.04.008

Bedny, M., Caramazza, A., Grossman, E., Pascual-Leone, A., & Saxe, R. (2008). Concepts are more than percepts: the case of action verbs. Journal of Neuroscience, 28(44), 11347–11353. https://doi.org/10.1523/JNEUROSCI.3039-08.2008

Bedny, M., Dravida, S., & Saxe, R. (2014). Shindigs, brunches, and rodeos: The neural basis of event words. Cognitive, Affective, & Behavioral Neuroscience, 14(3), 891–901. doi:10.3758/s13415-013-0217-z

Bemis, D. K., & Pylkkänen, L. (2011). Simple composition: A magnetoencephalography investigation into the comprehension of minimal linguistic phrases. Journal of Neuroscience, 31(8), 2801–2814. https://doi.org/10.1523/JNEUROSCI.5003-10.2011

Bemis, D. K., & Pylkkanen, L. (2013). Combination across domains: an MEG investigation into the relationship between mathematical, pictorial, and linguistic processing. Frontiers in psychology, 3, 583. https://doi.org/10.3389/fpsyg.2012.00583

Bemis, D. K., & Pylkkänen, L. (2013). Flexible composition: MEG evidence for the deployment of basic combinatorial linguistic mechanisms in response to task demands. PloS one, 8(9), e73949. https://doi.org/10.1371/journal.pone.0073949

Benjamini, Y., & Hochberg, Y. (1995). Controlling the false discovery rate: a practical and powerful approach to multiple testing. Journal of the Royal statistical society: series B (Methodological), 57(1), 289–300. https://doi.org/10.1111/j.2517-6161.1995.tb02031.x

Boylan, C., Trueswell, J. C., & Thompson-Schill, S. L. (2015). Compositionality and the angular gyrus: A multi-voxel similarity analysis of the semantic composition of nouns and verbs. Neuropsychologia, 78, 130–141. https://doi.org/10.1016/j.neuropsychologia.2015.10.007

Boylan, C., Trueswell, J. C., & Thompson-Schill, S. L. (2017). Relational vs. attributive interpretation of nominal compounds differentially engages angular gyrus and anterior temporal lobe. Brain and language, 169, 8–21. https://doi.org/10.1016/j.bandl.2017.01.008

Chan, A. M., Baker, J. M., Eskandar, E., Schomer, D., Ulbert, I., Marinkovic, K., Cash, S. S., & Halgren, E. (2011). First-pass selectivity for semantic categories in human anteroventral temporal lobe. Journal of Neuroscience, 31(49), 18119–18129. https://doi.org/10.1523/JNEUROSCI.3122-11.2011

Clarke, A., Taylor, K. I., & Tyler, L. K. (2011). The evolution of meaning: spatio-temporal dynamics of visual object recognition. Journal of cognitive neuroscience, 23(8), 1887–1899. https://doi.org/10.1162/jocn.2010.21544

Clarke, A., Taylor, K. I., Devereux, B., Randall, B., & Tyler, L. K. (2013). From perception to conception: how meaningful objects are processed over time. Cerebral Cortex, 23(1), 187–197. https://doi.org/10.1093/cercor/bhs002

Cohen, L., Lehéricy, S., Chochon, F., Lemer, C., Rivaud, S., & Dehaene, S. (2002). Language‐specific tuning of visual cortex? Functional properties of the Visual Word Form Area. Brain, 125(5), 1054–1069. https://doi.org/10.1093/brain/awf094

Coutanche, M. N., & Thompson-Schill, S. L. (2015). Creating concepts from converging features in human cortex. Cerebral cortex, 25(9), 2584–2593. https://doi.org/10.1093/cercor/bhu057

Dale, A. M., & Sereno, M. I. (1993). Improved localization of cortical activity by combining EEG and MEG with MRI cortical surface reconstruction: a linear approach. Journal of cognitive neuroscience, 5(2), 162–176. https://doi.org/10.1162/jocn.1993.5.2.162

Fischl, B., Liu, A., & Dale, A. M. (2001). Automated manifold surgery: constructing geometrically accurate and topologically correct models of the human cerebral cortex. IEEE transactions on medical imaging, 20(1), 70–80. DOI: 10.1109/42.906426

Gramfort, A., Luessi, M., Larson, E., Engemann, D. A., Strohmeier, D., Brodbeck, C., Goj, R., Jas, M., Brooks, T., Parkkonen, L., & Hämäläinen, M. (2013). MEG and EEG data analysis with MNE-Python. Frontiers in neuroscience, 7, 267. https://doi.org/10.3389/fnins.2013.00267

Dale, A. M., Fischl, B., & Sereno, M. I. (1999). Cortical surface-based analysis: I. Segmentation and surface reconstruction. Neuroimage, 9(2), 179–194. https://doi.org/10.1006/nimg.1998.0395

Davey, J., Cornelissen, P. L., Thompson, H. E., Sonkusare, S., Hallam, G., Smallwood, J., & Jefferies, E. (2015). Automatic and controlled semantic retrieval: TMS reveals distinct contributions of posterior middle temporal gyrus and angular gyrus. Journal of Neuroscience, 35(46), 15230–15239. https://doi.org/10.1523/JNEUROSCI.4705-14.2015

Davey, J., Thompson, H. E., Hallam, G., Karapanagiotidis, T., Murphy, C., De Caso, I., … & Jefferies, E. (2016). Exploring the role of the posterior middle temporal gyrus in semantic cognition: Integration of anterior temporal lobe with executive processes. Neuroimage, 137, 165–177. https://doi.org/10.1016/j.neuroimage.2016.05.051

Davies, M. (2009). The 385+ million word Corpus of Contemporary American English (1990– 2008+): Design, architecture, and linguistic insights. International journal of corpus linguistics, 14(2), 159–190. https://doi.org/10.1075/ijcl.14.2.02dav

Davis, M. H., Meunier, F., & Marslen-Wilson, W. D. (2004). Neural responses to morphological, syntactic, and semantic properties of single words: an fMRI study. Brain and language, 89(3), 439–449. https://doi.org/10.1016/S0093-934X(03)00471-1

de Zubicaray, G. I., Hansen, S., & McMahon, K. L. (2013). Differential processing of thematic and categorical conceptual relations in spoken word production. Journal of Experimental Psychology: General, 142(1), 131. https://doi.org/10.1037/a0028717

Dehaene, S., Pegado, F., Braga, L. W., Ventura, P., Nunes Filho, G., Jobert, A., Dehaene-Lambertz, G., Kolinsky, R., & Cohen, L. (2010). How learning to read changes the cortical networks for vision and language. science, 330(6009), 1359–1364. DOI: 10.1126/science.1194140

Desikan, R. S., Ségonne, F., Fischl, B., Quinn, B. T., Dickerson, B. C., Blacker, D., Buckner, R. L., Dale, A. M., Maguire, R.P., Hyman, B., T., Albert, M.S., & Killiany, R.J. (2006). An automated labeling system for subdividing the human cerebral cortex on MRI scans into gyral based regions of interest. Neuroimage, 31(3), 968–980. https://doi.org/10.1016/j.neuroimage.2006.01.021

Estes, Z. (2003). A tale of two similarities: Comparison and integration in conceptual combination. Cognitive Science, 27(6), 911–921. https://doi.org/10.1207/s15516709cog2706_4

Estes, Z., Golonka, S., & Jones, L. L. (2011). Thematic thinking: The apprehension and consequences of thematic relations. In Psychology of learning and motivation (Vol. 54, pp. 249–294). Academic Press. https://doi.org/10.1016/B978-0-12-385527-5.00008-5

Fedorenko, E., Hsieh, P. J., Nieto-Castañón, A., Whitfield-Gabrieli, S., & Kanwisher, N. (2010). New method for fMRI investigations of language: defining ROIs functionally in individual subjects. Journal of neurophysiology, 104(2), 1177–1194. https://doi.org/10.1152/jn.00032.2010

Fischl, B., Van Der Kouwe, A., Destrieux, C., Halgren, E., Ségonne, F., Salat, D. H., Busa, E., Seidman, L. J., Goldstein, J., Kennedy, D., Caviness, V., Makris, N., Rosen, B., & Dale, A. M. (2004). Automatically parcellating the human cerebral cortex. Cerebral cortex, 14(1), 11–22. https://doi.org/10.1093/cercor/bhg087

Fischl, B., Sereno, M. I., & Dale, A. M. (1999). Cortical surface-based analysis: II: inflation, flattening, and a surface-based coordinate system. Neuroimage, 9(2), 195–207. https://doi.org/10.1006/nimg.1998.0396

Flick, G., Oseki, Y., Kaczmarek, A. R., Al Kaabi, M., Marantz, A., & Pylkkänen, L. (2018). Building words and phrases in the left temporal lobe. Cortex, 106, 213–236. https://doi.org/10.1016/j.cortex.2018.06.004

Flick, G., & Pylkkänen, L. (2020). Isolating syntax in natural language: MEG evidence for an early contribution of left posterior temporal cortex. Cortex, 127, 42–57. https://doi.org/10.1016/j.cortex.2020.01.025

Frankland, S. M., & Greene, J. D. (2015). An architecture for encoding sentence meaning in left mid-superior temporal cortex. Proceedings of the National Academy of Sciences, 112(37), 11732–11737. https://doi.org/10.1073/pnas.1421236112

Frankland, S. M., & Greene, J. D. (2020). Two ways to build a thought: distinct forms of compositional semantic representation across brain regions. Cerebral Cortex, 30(6), 3838–3855. https://doi.org/10.1093/cercor/bhaa001

Gagné, C. L., & Shoben, E. J. (1997). Influence of thematic relations on the comprehension of modifier–noun combinations. Journal of experimental psychology: Learning, memory, and cognition, 23(1), 71. https://doi.org/10.1037/0278-7393.23.1.71

Glasser, M. F., Sotiropoulos, S. N., Wilson, J. A., Coalson, T. S., Fischl, B., Andersson, J. L., Xu, J., Jbabdi, S., Webster, M., Polimenni, J.R., Van Essen, D. C., & Jenkinson, M. (2013). The minimal preprocessing pipelines for the Human Connectome Project. Neuroimage, 80, 105–124. https://doi.org/10.1016/j.neuroimage.2013.04.127

Graves, W. W., Binder, J. R., Desai, R. H., Conant, L. L., & Seidenberg, M. S. (2010). Neural correlates of implicit and explicit combinatorial semantic processing. Neuroimage, 53(2), 638–646. https://doi.org/10.1016/j.neuroimage.2010.06.055

Gray, K. C., & Smith, E. E. (1995). The role of instance retrieval in understanding complex concepts. Memory & Cognition, 23(6), 665–674. https://doi.org/10.3758/BF03200920

Gwilliams, L., Lewis, G. A., & Marantz, A. (2016). Functional characterisation of letter-specific responses in time, space and current polarity using magnetoencephalography. Neuroimage, 132, 320–333. https://doi.org/10.1016/j.neuroimage.2016.02.057

Hallam, G. P., Thompson, H. E., Hymers, M., Millman, R. E., Rodd, J. M., Ralph, M. A. L., Smallwood, J., & Jefferies, E. (2018). Task-based and resting-state fMRI reveal compensatory network changes following damage to left inferior frontal gyrus. Cortex, 99, 150–165. https://doi.org/10.1016/j.cortex.2017.10.004

Hallam, G. P., Whitney, C., Hymers, M., Gouws, A. D., & Jefferies, E. (2016). Charting the effects of TMS with fMRI: Modulation of cortical recruitment within the distributed network supporting semantic control. Neuropsychologia, 93, 40–52. https://doi.org/10.1016/j.neuropsychologia.2016.09.012

Hämäläinen, M., Hari, R., Ilmoniemi, R. J., Knuutila, J., & Lounasmaa, O. V. (1993). Magnetoencephalography—theory, instrumentation, and applications to noninvasive studies of the working human brain. Reviews of modern Physics, 65(2), 413. https://doi.org/10.1103/RevModPhys.65.413

Hauk, O., Wakeman, D. G., & Henson, R. (2011). Comparison of noise-normalized minimum norm estimates for MEG analysis using multiple resolution metrics. Neuroimage, 54(3), 1966–1974. https://doi.org/10.1016/j.neuroimage.2010.09.053

Hauk, O., Stenroos, M., & Treder, M. (2019). EEG/MEG source estimation and spatial filtering: the linear toolkit. In Supek S., Aine C. (eds) Magnetoencephalography. Spring, Cham. https://doi.org/10.1007/978-3-030-00087-5_85

Hillebrand, A., & Barnes, G. R. (2002). A quantitative assessment of the sensitivity of whole-head MEG to activity in the adult human cortex. Neuroimage, 16(3), 638–650. https://doi.org/10.1006/nimg.2002.1102

Humphreys, G. F., & Lambon Ralph, M. A. (2015). Fusion and fission of cognitive functions in the human parietal cortex. Cerebral Cortex, 25(10), 3547–3560. https://doi.org/10.1093/cercor/bhu198

Jackson, R. L. (2020). The Neural Correlates of Semantic Control Revisited. NeuroImage, 117444. https://doi.org/10.1016/j.neuroimage.2020.117444

Jefferies, E. (2013). The neural basis of semantic cognition: converging evidence from neuropsychology, neuroimaging and TMS. Cortex, 49(3), 611–625. https://doi.org/10.1016/j.cortex.2012.10.008

Jefferies, E., Thompson, H., Cornelissen, P., & Smallwood, J. (2020). The neurocognitive basis of knowledge about object identity and events: dissociations reflect opposing effects of semantic coherence and control. Philosophical Transactions of the Royal Society B, 375(1791), 20190300. https://doi.org/10.1098/rstb.2019.0300

Kable, J. W., Kan, I. P., Wilson, A., Thompson-Schill, S. L., & Chatterjee, A. (2005). Conceptual representations of action in the lateral temporal cortex. Journal of cognitive neuroscience, 17(12), 1855–1870. https://doi.org/10.1162/089892905775008625

Kable, J. W., Lease-Spellmeyer, J., & Chatterjee, A. (2002). Neural substrates of action event knowledge. Journal of Cognitive Neuroscience, 14(5), 795–805. https://doi.org/10.1162/08989290260138681

Kalénine, S., & Buxbaum, L. J. (2016). Thematic knowledge, artifact concepts, and the left posterior temporal lobe: Where action and object semantics converge. Cortex, 82, 164–178. https://doi.org/10.1016/j.cortex.2016.06.008

Kalénine, S., Mirman, D., Middleton, E. L., & Buxbaum, L. J. (2012). Temporal dynamics of activation of thematic and functional knowledge during conceptual processing of manipulable artifacts. Journal of Experimental Psychology: Learning, Memory, and Cognition, 38(5), 1274. https://doi.org/10.1037/a0027626

Kemmerer, D., Rudrauf, D., Manzel, K., & Tranel, D. (2012). Behavioral patterns and lesion sites associated with impaired processing of lexical and conceptual knowledge of actions. Cortex, 48(7), 826–848. https://doi.org/10.1016/j.cortex.2010.11.001

Kim, S., & Pylkkänen, L. (2019). Composition of event concepts: Evidence for distinct roles for the left and right anterior temporal lobes. Brain and language, 188, 18–27. https://doi.org/10.1016/j.bandl.2018.11.003

Lambon Ralph, M. A., Jefferies, E., Patterson, K., & Rogers, T. T. (2017). The neural and computational bases of semantic cognition. Nature Reviews Neuroscience, 18(1), 42. https://doi.org/10.1038/nrn.2016.150

Levi, J. N. (1978). The syntax and semantics of complex nominals. New York: Academic Press

Lewis, G., Solomyak, O., & Marantz, A. (2011). The neural basis of obligatory decomposition of suffixed words. Brain and language, 118(3), 118–127. https://doi.org/10.1016/j.bandl.2011.04.004

Lin, F. H., Belliveau, J. W., Dale, A. M., & Hämäläinen, M. S. (2006). Distributed current estimates using cortical orientation constraints. Human brain mapping, 27(1), 1–13. https://doi.org/10.1002/hbm.20155

Liu, A. K., Dale, A. M., & Belliveau, J. W. (2002). Monte Carlo simulation studies of EEG and MEG localization accuracy. Human brain mapping, 16(1), 47–62. https://doi.org/10.1002/hbm.10024

Margulies, D. S., Ghosh, S. S., Goulas, A., Falkiewicz, M., Huntenburg, J. M., Langs, G., Bezgin, G., Eickhoff, S. B., Castellanos, F.X., Petrides, M., Jefferies, E., & Smallwood, J. (2016). Situating the default-mode network along a principal gradient of macroscale cortical organization. Proceedings of the National Academy of Sciences, 113(44), 12574–12579. https://doi.org/10.1073/pnas.1608282113

Maris, E., & Oostenveld, R. (2007). Nonparametric statistical testing of EEG-and MEG-data. Journal of neuroscience methods, 164(1), 177–190. https://doi.org/10.1016/j.jneumeth.2007.03.024

Martin, A., Haxby, J. V., Lalonde, F. M., Wiggs, C. L., & Ungerleider, L. G. (1995). Discrete cortical regions associated with knowledge of color and knowledge of action. Science, 270(5233), 102–105. DOI: 10.1126/science.270.5233.102

Matchin, W., Brodbeck, C., Hammerly, C., & Lau, E. (2019). The temporal dynamics of structure and content in sentence comprehension: Evidence from fMRI‐constrained MEG. Human brain mapping, 40(2), 663–678. https://doi.org/10.1002/hbm.24403

Matchin, W., Hammerly, C., & Lau, E. (2017). The role of the IFG and pSTS in syntactic prediction: Evidence from a parametric study of hierarchical structure in fMRI. Cortex, 88, 106–123. https://doi.org/10.1016/j.cortex.2016.12.010

Mirman, D., & Graziano, K. M. (2012a). Individual differences in the strength of taxonomic versus thematic relations. Journal of experimental psychology: General, 141(4), 601. https://doi.org/10.1037/a0026451

Mirman, D., & Graziano, K. M. (2012b). Damage to temporo-parietal cortex decreases incidental activation of thematic relations during spoken word comprehension. Neuropsychologia, 50(8), 1990–1997. https://doi.org/10.1016/j.neuropsychologia.2012.04.024

Mirman, D., Landrigan, J. F., & Britt, A. E. (2017). Taxonomic and thematic semantic systems. Psychological bulletin, 143(5), 499. https://doi.org/10.1037/bul0000092

Molins, A., Stufflebeam, S. M., Brown, E. N., & Hämäläinen, M. S. (2008). Quantification of the benefit from integrating MEG and EEG data in minimum ℓ2-norm estimation. Neuroimage, 42(3), 1069–1077. https://doi.org/10.1016/j.neuroimage.2008.05.064

Moss, H. E., Rodd, J. M., Stamatakis, E. A., Bright, P., & Tyler, L. K. (2005). Anteromedial temporal cortex supports fine-grained differentiation among objects. Cerebral cortex, 15(5), 616–627. https://doi.org/10.1093/cercor/bhh163

Murphy, G. L. (1988). Comprehending complex concepts. Cognitive science, 12(4), 529–562. https://doi.org/10.1207/s15516709cog1204_2

Murphy, G. L. (1990). Noun phrase interpretation and conceptual combination. Journal of memory and language, 29(3), 259–288. https://doi.org/10.1016/0749-596X(90)90001-G

Neophytou, K., Manouilidou, C., Stockall, L., & Marantz, A. (2018). Syntactic and semantic restrictions on morphological recomposition: MEG evidence from Greek. Brain and language, 183, 11–20. https://doi.org/10.1016/j.bandl.2018.05.003

Noonan, K. A., Jefferies, E., Visser, M., & Lambon Ralph, M. A. (2013). Going beyond inferior prefrontal involvement in semantic control: evidence for the additional contribution of dorsal angular gyrus and posterior middle temporal cortex. Journal of cognitive neuroscience, 25(11), 1824–1850. https://doi.org/10.1162/jocn_a_00442

Okada, Y. C., Wu, J., & Kyuhou, S. (1997). Genesis of MEG signals in a mammalian CNS structure. Electroencephalography and clinical neurophysiology, 103(4), 474–485. https://doi.org/10.1016/S0013-4694(97)00043-6

Pallier, C., Devauchelle, A. D., & Dehaene, S. (2011). Cortical representation of the constituent structure of sentences. Proceedings of the National Academy of Sciences, 108(6), 2522–2527. https://doi.org/10.1073/pnas.1018711108

Patterson, K., Nestor, P. J., & Rogers, T. T. (2007). Where do you know what you know? The representation of semantic knowledge in the human brain. Nature reviews neuroscience, 8(12), 976–987. https://doi.org/10.1038/nrn2277

Price, A. R., Bonner, M. F., Peelle, J. E., & Grossman, M. (2015). Converging evidence for the neuroanatomic basis of combinatorial semantics in the angular gyrus. Journal of Neuroscience, 35(7), 3276–3284. https://doi.org/10.1523/JNEUROSCI.3446-14.2015

Price, A. R., Peelle, J. E., Bonner, M. F., Grossman, M., & Hamilton, R. H. (2016). Causal evidence for a mechanism of semantic integration in the angular gyrus as revealed by high-definition transcranial direct current stimulation. Journal of Neuroscience, 36(13), 3829–3838. https://doi.org/10.1523/JNEUROSCI.3120-15.2016

Pylkkänen, L. (2020). Neural basis of basic composition: what we have learned from the red– boat studies and their extensions. Philosophical Transactions of the Royal Society B, 375(1791), 20190299. https://doi.org/10.1098/rstb.2019.0299

Pylkkänen, L., Stringfellow, A., & Marantz, A. (2002). Neuromagnetic evidence for the timing of lexical activation: An MEG component sensitive to phonotactic probability but not to neighborhood density. Brain and language, 81(1-3), 666–678. https://doi.org/10.1006/brln.2001.2555

Rodd, J. M., Longe, O. A., Randall, B., & Tyler, L. K. (2010). The functional organisation of the fronto-temporal language system: evidence from syntactic and semantic ambiguity. Neuropsychologia, 48(5), 1324–1335. https://doi.org/10.1016/j.neuropsychologia.2009.12.035

Rogalsky, C., LaCroix, A. N., Chen, K. H., Anderson, S. W., Damasio, H., Love, T., & Hickok, G. (2018). The neurobiology of agrammatic sentence comprehension: a lesion study. Journal of cognitive neuroscience, 30(2), 234–255. https://doi.org/10.1162/jocn_a_01200

Sassenhagen, J., & Draschkow, D. (2019). Cluster‐based permutation tests of MEG/EEG data do not establish significance of effect latency or location. Psychophysiology, 56(6), e13335. https://doi.org/10.1111/psyp.13335

Schmidtke, D., Gagné, C. L., Kuperman, V., & Spalding, T. L. (2018). Language experience shapes relational knowledge of compound words. Psychonomic bulletin & review, 25(4), 1468–1487. https://doi.org/10.3758/s13423-018-1478-x

Schwartz, M. F., Kimberg, D. Y., Walker, G. M., Brecher, A., Faseyitan, O. K., Dell, G. S., … & Coslett, H. B. (2011). Neuroanatomical dissociation for taxonomic and thematic knowledge in the human brain. Proceedings of the National Academy of Sciences, 108(20), 8520–8524. https://doi.org/10.1073/pnas.1014935108

Smith, S. M., & Nichols, T. E. (2009). Threshold-free cluster enhancement: addressing problems of smoothing, threshold dependence and localisation in cluster inference. Neuroimage, 44(1), 83–98. https://doi.org/10.1016/j.neuroimage.2008.03.061

Snijders, T. M., Vosse, T., Kempen, G., Van Berkum, J. J., Petersson, K. M., & Hagoort, P. (2009). Retrieval and unification of syntactic structure in sentence comprehension: an fMRI study using word-category ambiguity. Cerebral cortex, 19(7), 1493–1503. https://doi.org/10.1093/cercor/bhn187

Solomyak, O., & Marantz, A. (2010). Evidence for early morphological decomposition in visual word recognition. Journal of Cognitive Neuroscience, 22(9), 2042–2057. https://doi.org/10.1162/jocn.2009.21296

Tarkiainen, A., Helenius, P., Hansen, P. C., Cornelissen, P. L., & Salmelin, R. (1999). Dynamics of letter string perception in the human occipitotemporal cortex. Brain, 122(11), 2119–2132. https://doi.org/10.1093/brain/122.11.2119

Taylor, J. S. H., Davis, M. H., & Rastle, K. (2019). Mapping visual symbols onto spoken language along the ventral visual stream. Proceedings of the National Academy of Sciences, 116(36), 17723–17728. https://doi.org/10.1073/pnas.1818575116

Teige, C., Mollo, G., Millman, R., Savill, N., Smallwood, J., Cornelissen, P. L., & Jefferies, E. (2018). Dynamic semantic cognition: Characterising coherent and controlled conceptual retrieval through time using magnetoencephalography and chronometric transcranial magnetic stimulation. Cortex, 103, 329–349. https://doi.org/10.1016/j.cortex.2018.03.024

Teige, C., Cornelissen, P. L., Mollo, G., Alam, T. R. D. J. G., McCarty, K., Smallwood, J., & Jefferies, E. (2019). Dissociations in semantic cognition: Oscillatory evidence for opposing effects of semantic control and type of semantic relation in anterior and posterior temporal cortex. cortex, 120, 308–325. https://doi.org/10.1016/j.cortex.2019.07.002

Thompson, H., Davey, J., Hoffman, P., Hallam, G., Kosinski, R., Howkins, S., Wooffindin, E., Gabbitas, R., & Jefferies, E. (2017). Semantic control deficits impair understanding of thematic relationships more than object identity. Neuropsychologia, 104, 113–125. https://doi.org/10.1016/j.neuropsychologia.2017.08.013

Tyler, L. K., Cheung, T. P., Devereux, B. J., & Clarke, A. (2013). Syntactic computations in the language network: characterizing dynamic network properties using representational similarity analysis. Frontiers in psychology, 4, 271. https://doi.org/10.3389/fpsyg.2013.00271

Tyler, L. K., Stamatakis, E. A., Bright, P., Acres, K., Abdallah, S., Rodd, J. M., & Moss, H. E. (2004). Processing objects at different levels of specificity. Journal of cognitive neuroscience, 16(3), 351–362. https://doi.org/10.1162/089892904322926692

Van Essen, D. C. (2005). A population-average, landmark-and surface-based (PALS) atlas of human cerebral cortex. Neuroimage, 28(3), 635–662. https://doi.org/10.1016/j.neuroimage.2005.06.058

Vinckier, F., Dehaene, S., Jobert, A., Dubus, J. P., Sigman, M., & Cohen, L. (2007). Hierarchical coding of letter strings in the ventral stream: dissecting the inner organization of the visual word-form system. Neuron, 55(1), 143–156. https://doi.org/10.1016/j.neuron.2007.05.031

Wang, J., Deng, Y., & Booth, J. R. (2019). Automatic semantic influence on early visual word recognition in the ventral occipito-temporal cortex. Neuropsychologia, 133, 107188. https://doi.org/10.1016/j.neuropsychologia.2019.107188

Weisberg, J., Van Turennout, M., & Martin, A. (2007). A neural system for learning about object function. Cerebral Cortex, 17(3), 513–521. https://doi.org/10.1093/cercor/bhj176

Westerlund, M., & Pylkkänen, L. (2014). The role of the left anterior temporal lobe in semantic composition vs. semantic memory. Neuropsychologia, 57, 59–70. https://doi.org/10.1016/j.neuropsychologia.2014.03.001

Whitney, C., Kirk, M., O’Sullivan, J., Lambon Ralph, M. A., & Jefferies, E. (2011). The neural organization of semantic control: TMS evidence for a distributed network in left inferior frontal and posterior middle temporal gyrus. Cerebral Cortex, 21(5), 1066–1075. https://doi.org/10.1093/cercor/bhq180

Williams, A., Reddigari, S., & Pylkkänen, L. (2017). Early sensitivity of left perisylvian cortex to relationality in nouns and verbs. Neuropsychologia, 100, 131–143. https://doi.org/10.1016/j.neuropsychologia.2017.04.029

Wisniewski, E. J., & Love, B. C. (1998). Relations versus properties in conceptual combination. Journal of memory and language, 38(2), 177–202. https://doi.org/10.1006/jmla.1997.2550

Wisniewski, E. J. (1996). Construal and similarity in conceptual combination. Journal of Memory and Language, 35(3), 434–453. https://doi.org/10.1006/jmla.1996.0024

Woolnough, O., Donos, C., Rollo, P. S., Forseth, K. J., Lakretz, Y., Crone, N. E., Fischer-Baum, S., Dehaene, S., & Tandon, N. (2020). Spatiotemporal dynamics of orthographic and lexical processing in the ventral visual pathway. Nature human behaviour, 10.1038/s41562-020-00982-w. Advance online publication. https://doi.org/10.1038/s41562-020-00982-w

Yu, X., Bi, Y., Han, Z., Zhu, C., & Law, S. P. (2012). Neural correlates of comprehension and production of nouns and verbs in Chinese. Brain and language, 122(2), 126–131. https://doi.org/10.1016/j.bandl.2012.05.002

Yu, X., Law, S. P., Han, Z., Zhu, C., & Bi, Y. (2011). Dissociative neural correlates of semantic processing of nouns and verbs in Chinese—A language with minimal inflectional morphology. NeuroImage, 58(3), 912–922. https://doi.org/10.1016/j.neuroimage.2011.06.039

Zhang, L., & Pylkkänen, L. (2015). The interplay of composition and concept specificity in the left anterior temporal lobe: An MEG study. NeuroImage, 111, 228–240. https://doi.org/10.1016/j.neuroimage.2015.02.028

Ziegler, J., & Pylkkänen, L. (2016). Scalar adjectives and the temporal unfolding of semantic composition: An MEG investigation. Neuropsychologia, 89, 161–171. https://doi.org/10.1016/j.neuropsychologia.2016.06.010

