## Supplementary Materials for "From letters to composed concepts: A magnetoencephalography study of reading"

##### Stimulus Norming Studies

To create the materials used in Experiment 2 (noun-noun combinations) we conducted two stimulus norming studies on Amazon Mechanical Turk (AMT). All participants were self-declared native English speakers, with Internet Protocol addresses located in the United States of America. The first norming study asked participants to rate on a 7-point Likert scale their overall familiarity with the meanings of noun-noun compounds (7 = extremely familiar, 1 = never considered before). Each participant was presented with a list of 42 noun-noun compounds, containing 13 of the experimental stimuli (6 of one condition, 7 of the other). No constituent noun was repeated within the list for a given participant. The remaining thirty compounds in each list were a constant set of fillers, selected from the stimuli of Graves et al., (2013) to span the distribution of meaningfulness ratings collected in that work, with 7, 7, 8, and 8 compounds from the first, second, third, and fourth quartiles, respectively. After removing participants who failed to complete all ratings, between 21 and 30 participants (mean = 26.47) responded to each noun-noun combination. The mean familiarity ratings were 4.750 (sd = 1.021) and 4.838 (sd = 1.351) in the Material and Contents conditions, respectively, indicating similar familiarity between the two conditions and an overall high level of familiarity with the compounds. A t-test of the difference in familiarity scores across conditions did not reveal a statistically significance difference ($t(41) = 0.638$, $p < 0.73$).

The second AMT norming study was designed to confirm that participants interpreted the noun-noun combinations with the intended semantic relation between the constituents (i.e., that *leather bag* is a bag made of leather rather than a bag that holds leather). This was done by presenting the combinations to each AMT participant and asking them to select from ten choices the best paraphrase that matched their interpretation. Each combination was presented in the following frame sentence: A [modifier] [head] is a [head] _____ [modifier]. For example: “a steel cabinet is a cabinet ____ steel”. A drop-down box then provided participants with the following response options that could be used to paraphrase the combination’s meaning: “about”, “during”, “for”, “found in/on/near”, “made from”, “made of”, “that causes”, “that looks like”, “that makes”, and “No acceptable answer”. Participants were instructed to select the last option if none of the other choices made intuitive sense to them. The relation choices were motivated by previous psycholinguistic analyses of English noun-noun compounding (Levi, 1979). If a modifier instantiated a count noun, it was preceded by the indefinite determiner after the blank space. All other nouns were presented without a determiner in the final position of the frame. Each participant was given a list of 41 compounds. As in the first norming study, 13 of these were experimental stimuli (6 and 7 from each condition) and the remaining 28 were taken from the quartiles of the Graves et al., (2013) materials. The choice of the “for” response (i.e., *trophy cabinet is a cabinet for a trophy*) was coded as indicating a Contents relationship, while the choice of either “made of” or “made from” was coded as indicating a Material relationship. A measure of relation dominance was then defined as the proportion of times that participants selected the intended relations. After removing participants who did not complete the task, between 26 and 32 individuals (mean = 29.17) judged each stimulus. The results indicated that the two conditions were very similar in their dominance scores: Material = 0.918 (sd=0.092) and Contents = 0.899 (0.060), and that the majority of participants interpreted the stimuli in the intended manner.

**References:** Graves, W. W., Binder, J. R., & Seidenberg, M. S. (2013). Noun–noun combination: Meaningfulness ratings and lexical statistics for 2,160 word pairs. *Behavior research methods*, *45*(2), 463-469.

### Supplementary Figures

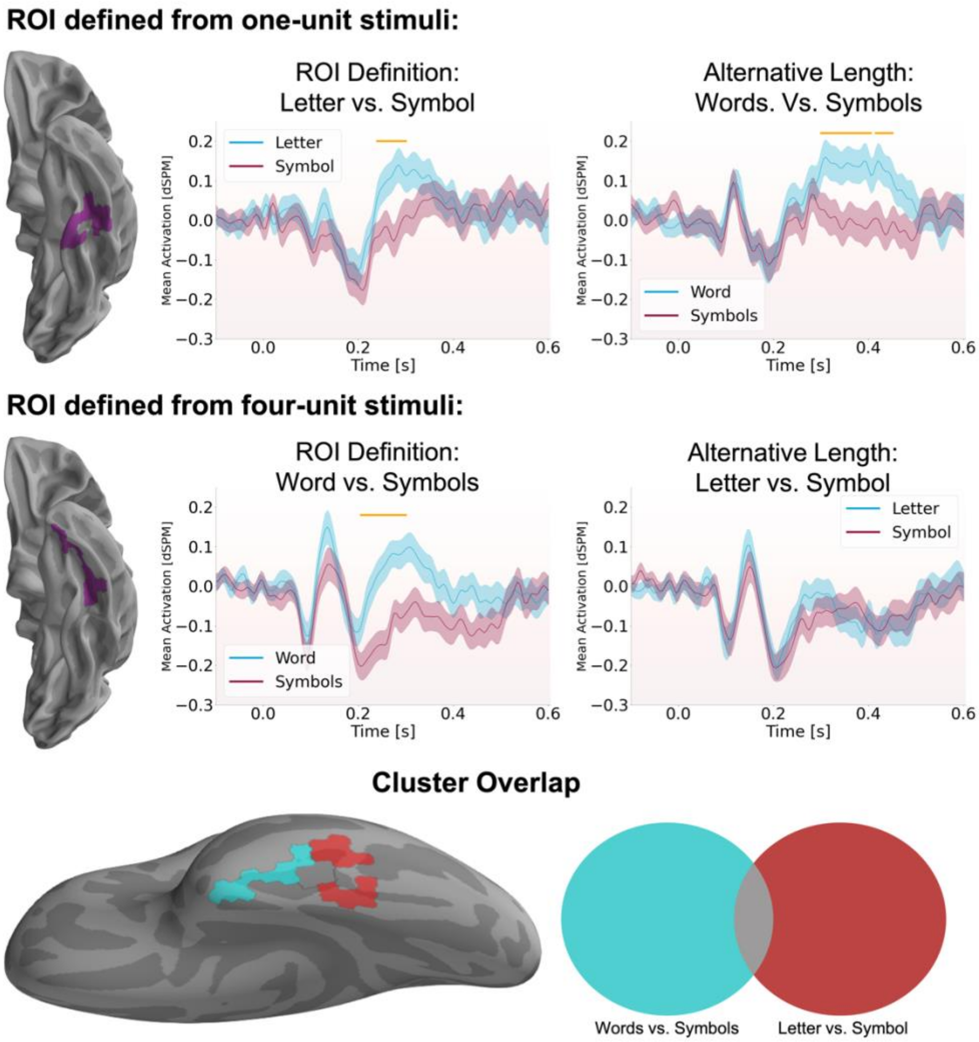


**Supplementary Figure 1.** Examination of ventral responses discriminating individual letters vs. individual symbols (top) and words vs. four-unit symbol strings (middle). The left panel in each of the first two rows displays the largest cluster identified in the analysis using the corresponding length stimuli. The right panel shows the response, in this cluster, to the alternative length stimulus contrast. The cluster identified from the contrast of letters vs. one symbol also discriminated between words and four symbols, although at a later time relative to stimulus onset (top-right: 100-600 ms test window, significant clusters: 300-405 ms, uncorrected $p = 0.0075$; 415-450 ms, uncorrected $p = 0.0042$). On the other hand, the cluster identified from the contrast of words vs. four symbols, which was located more anteriorly along the ventral surface, did not discriminate between individual letters and symbols. The bottom panel shows the spatial overlap of the two and highlights the more anterior extent of the cluster identified from the word vs. four-symbol contrast.

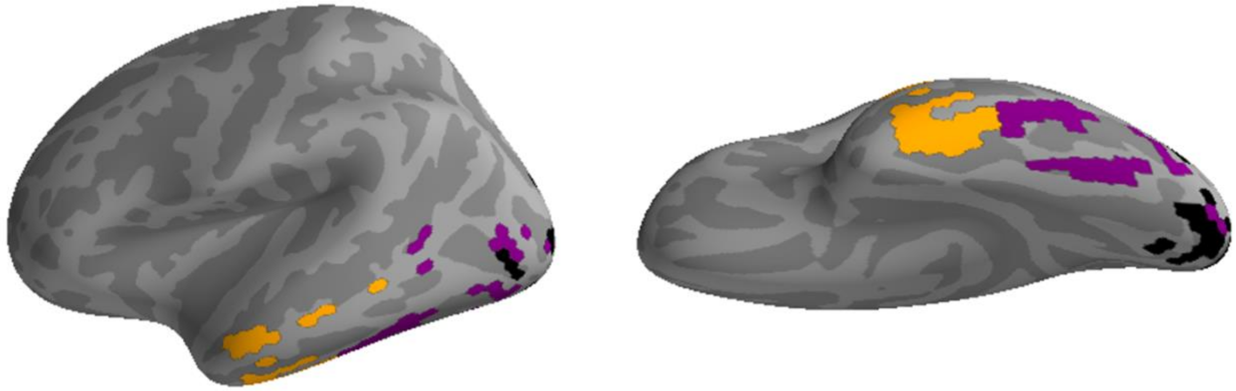

**Supplementary Figure 2.** Crosstalk functions for three clusters found in the analysis of the letter-string response localizer: (1) The lateral occipital cluster that distinguished high and low visual noise stimuli (black); (2) the posterior fusiform cluster that distinguished high and low visual noise stimuli (purple); and (3) the anterior fusiform and inferior temporal lobe that distinguished between four-letter words and four-unit symbol strings. All CTFs were masked using a half maximum threshold on the absolute values.

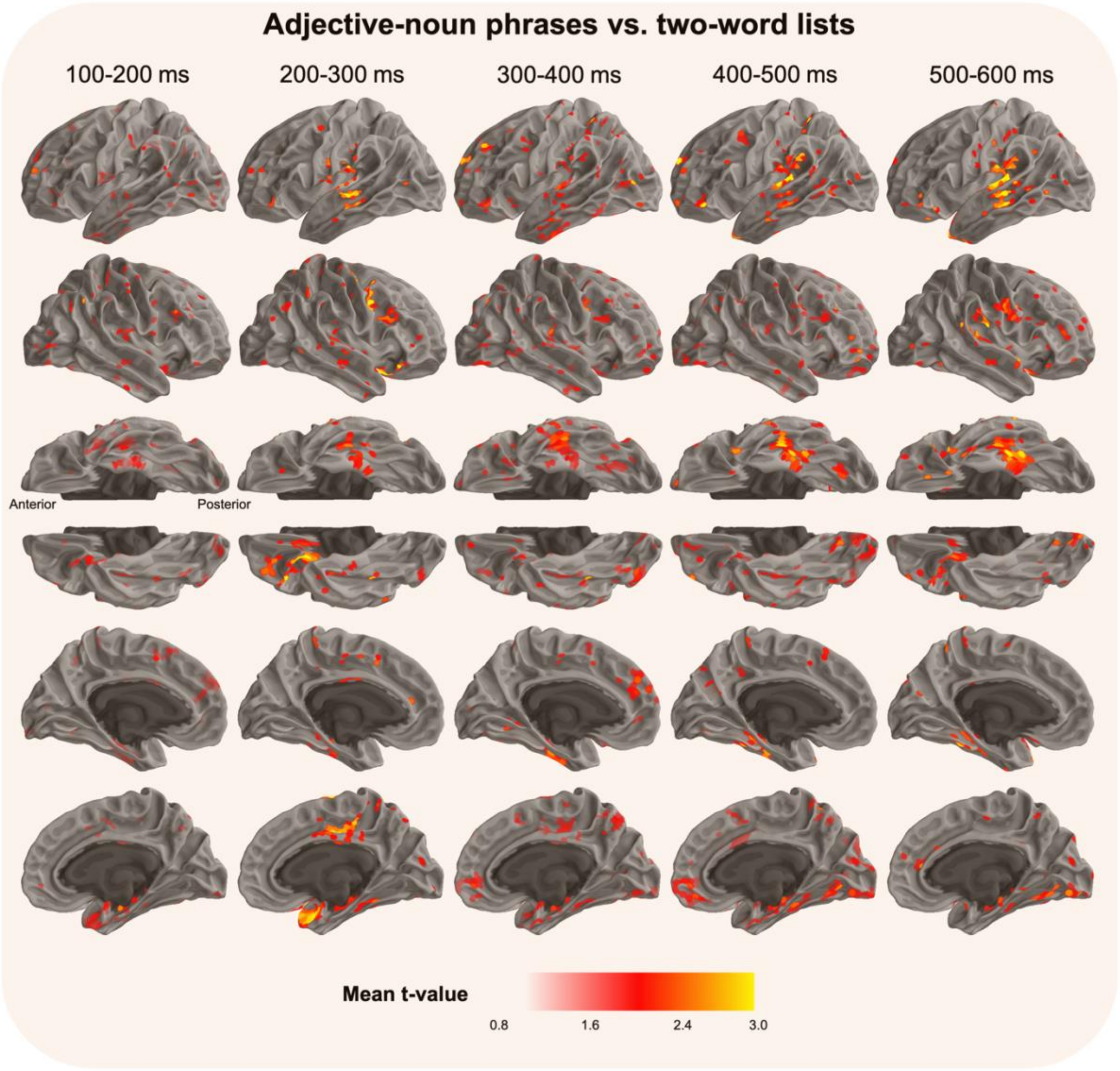


**Supplementary Figure 3.** Experiment 1. Whole brain statistical contrast of two-word phrases (red boat) vs. two-word lists (cup, boat). Each column shows the mean absolute t-value across the indicated time window. All times are relative to the onset of the second word of each trial (i.e., the phrasal head).

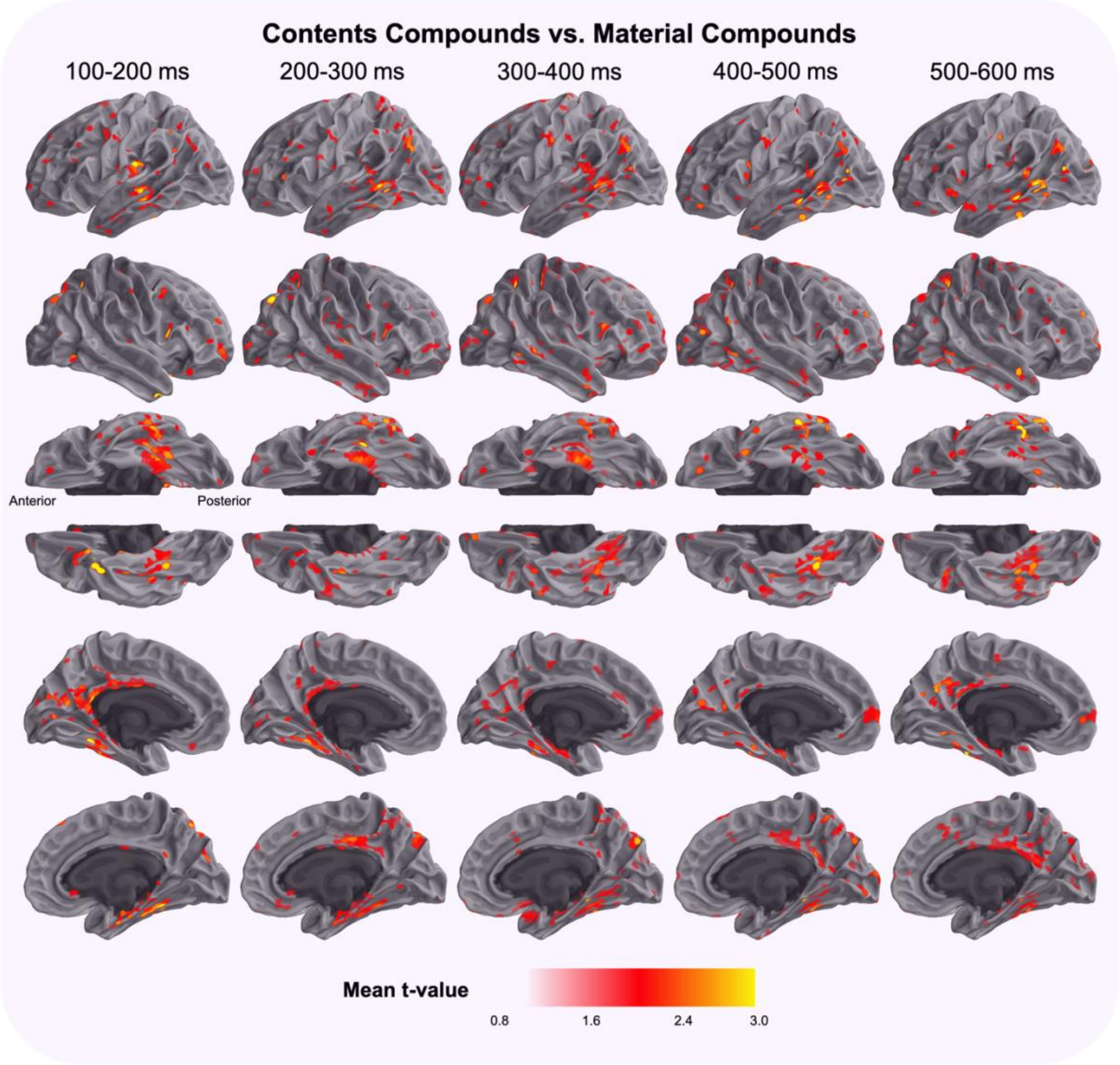


**Supplementary Figure 4.** Experiment 2. Whole brain statistical contrast of Contents Compounds (trophy cabinet) vs. Material Compounds (metal cabinet). Each column shows the mean absolute t-value across the indicated time window. All times are relative to the onset of the second word of each trial (i.e., the phrasal head).

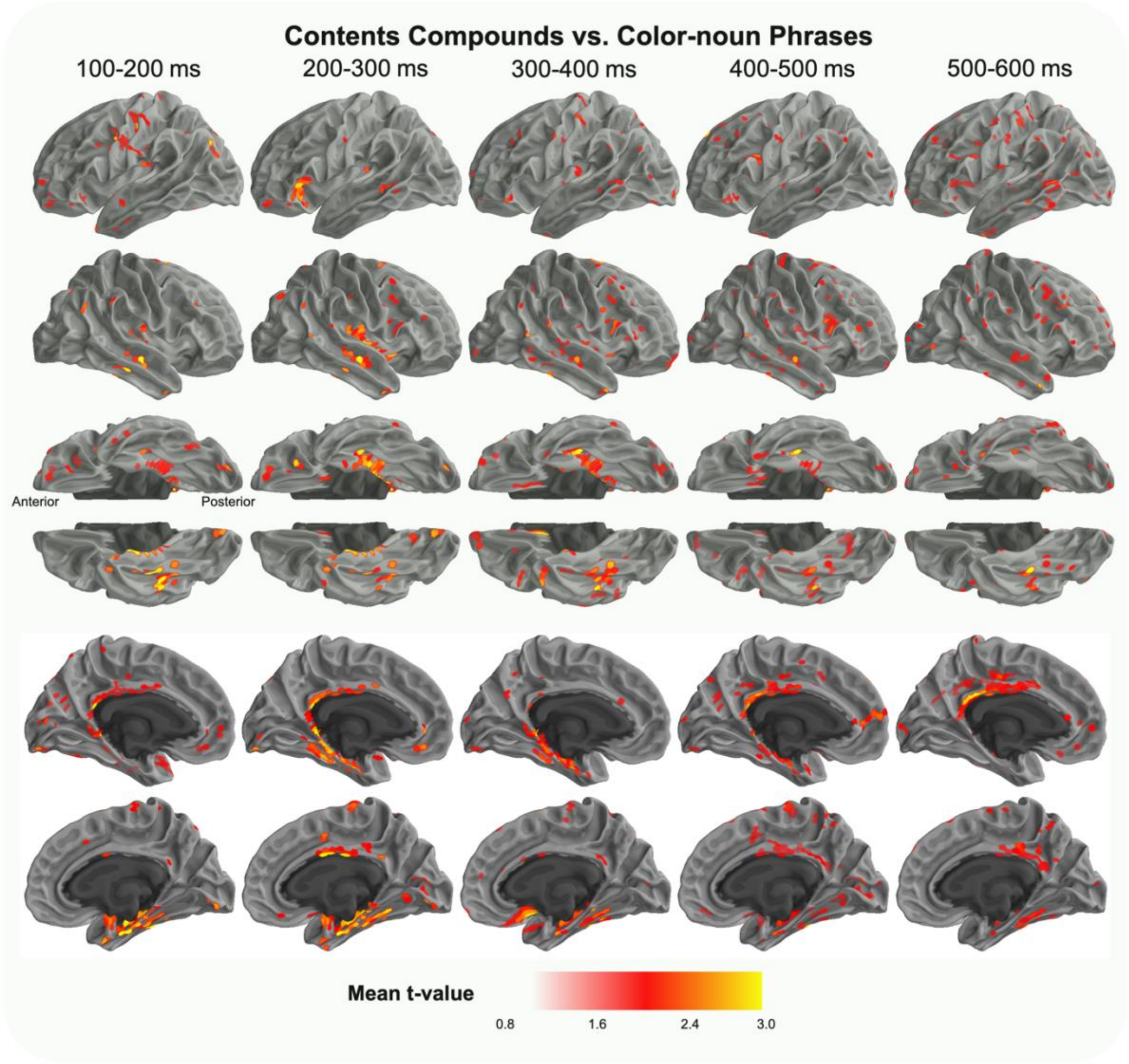


**Supplementary Figure 5.** Experiment 2. Whole brain statistical contrast of Contents Compounds (trophy cabinet) vs. Color-noun phrases (green cabinet). Each column shows the mean absolute t-value across the indicated time window. All times are relative to the onset of the second word of each trial (i.e., the phrasal head).

#### Supplementary Tables

Supplementary Table 1. All spatiotemporal clusters found in analyses of the letter-string response localizer. The indicated location of each cluster is the label of the automated surface parcellation that contained the greatest number of that cluster's constituent sources.

| Test Name | Cluster Duration | Location | Number of Sources | p-value |
|---|---|---|---|---|
| Noise Level | 130-180 ms | Lateral occipital | 47 | < 0.0001 |
| | 130-180 ms | Fusiform | 14 | < 0.0001 |
| | 130-165 ms | Lingual gyrus | 7 | 0.0095 |
| | 150-155 ms | Fusiform | 1 | 0.0460 |
| | 130-160 ms | Lateral occipital | 2 | 0.0180 |
| | 140-155 ms | Fusiform | 2 | 0.0317 |
| | 145-160 ms | Fusiform | 2 | 0.0381 |
| | 130-155 ms | Fusiform | 2 | 0.0229 |
| | 165-180 ms | fusiform | 1 | 0.0318 |
| String Type | 210-300 ms | Inferior temporal | 21 | 0.0004 |
| | 235-300 ms | Middle temporal | 2 | 0.0035 |
| | 290-300 ms | Fusiform | 1 | 0.0129 |
| String Type (only words vs. length-matched symbols) | 205-300 ms | Fusiform | 15 | 0.0008 |
| | 230-300 ms | Middle temporal | 1 | 0.0161 |
| | 245-265 ms | Inferior temporal | 3 | 0.0276 |
| | 285-300 ms | Inferior temporal | 2 | 0.0388 |
| | 215-300 ms | Fusiform | 1 | 0.0142 |

Supplementary Table 2: The complete set of stimuli used in the procedure of Experiment 2.

| Material Modifier | Contents Modifier | Color Adjective | Head |
|---|---|---|---|
| leather | garbage | white | bag |
| straw | laundry | black | bag |
| canvas | mail | brown | bag |
| metal | bread | green | barrel |
| oak | toy | red | barrel |
| clay | whisky | grey | barrel |
| metal | bread | green | basket |
| plastic | laundry | maroon | basket |
| straw | wine | black | basket |
| metal | bread | green | bin |
| aluminum | garbage | violet | bin |
| canvas | vegetable | brown | bin |
| plastic | shampoo | maroon | bottle |
| porcelain | soda | orange | bottle |
| tin | wine | blue | bottle |
| metal | bread | green | box |
| copper | jewelry | beige | box |
| oak | toy | red | box |
| metal | alcohol | green | bucket |
| titanium | bread | yellow | bucket |
| steel | champagne | purple | bucket |
| aluminum | alcohol | violet | cabinet |
| cedar | trophy | pink | cabinet |
| steel | vegetable | purple | cabinet |
| copper | jewelry | beige | chest |
| oak | toy | red | chest |
| cedar | trophy | pink | chest |
| plastic | champagne | maroon | crate |
| aluminum | vegetable | violet | crate |
| titanium | wine | yellow | crate |
| clay | shampoo | grey | cup |
| porcelain | soda | orange | cup |
| tin | whisky | blue | cup |
| plastic | soda | maroon | mug |
| clay | whisky | grey | mug |
| tin | wine | blue | mug |
| leather | garbage | white | sack |
| straw | laundry | black | sack |
| canvas | mail | brown | sack |
| copper | alcohol | beige | vault |
| titanium | champagne | yellow | vault |
| steel | jewelry | purple | vault |
